# Modeling cerebral development *in vitro* with L-*MYC*-immortalized human neural stem cell-derived organoids

**DOI:** 10.1101/2025.02.12.637976

**Authors:** Alejandra Velazquez Ojeda, Dina Awabdeh, Blake Brewster, Russell Rockne, Denis O’Meally, Hongwei Holly Yin, Nadia Carlesso, Christine E. Brown, Margarita Gutova, Michael E. Barish

**Affiliations:** Departments of Stem Cell Biology & Regenerative Medicine, Beckman Research Institute and National Medical Center, City of Hope, Duarte CA 91010; Departments of Computational & Quantitative Medicine, Beckman Research Institute and National Medical Center, City of Hope, Duarte CA 91010; Departments of Diabetes & Cancer Discovery, Beckman Research Institute and National Medical Center, City of Hope, Duarte CA 91010; Departments of Cancer Biology & Molecular Medicine, Beckman Research Institute and National Medical Center, City of Hope, Duarte CA 91010; Departments of Hematology & Hematopoietic Cell Transplantation, Beckman Research Institute and National Medical Center, City of Hope, Duarte CA 91010

**Keywords:** Human neural stem cells, LMNSC01 cells, NSC-derived organoids, neural lineage differentiation

## Abstract

A promising advance for *ex vivo* studies of human brain development and formulation of therapeutic strategies has been the adoption of brain organoids that, to a greater extent than monolayer or spheroid cultures, recapitulate to varying extents the patterns of tissue development and cell differentiation of human brain. Previously, such studies been hampered by limited access to relevant human tissue, inadequate human *in vitro* models, and the necessity of using rodent models that imperfectly reproduce human brain physiology. Here we present a novel organoid-based research platform utilizing L-*MYC*-immortalized human fetal neural stem cells (LMNSC01) grown in a physiological 4% oxygen environment.

We visualized developmental processes in LMNSC01 brain organoids for over 120 days *in vitro* by immunofluorescence and NanoString gene expression profiling. Gene expression patterns revealed by NanoString profiling were quantitatively compared to those occurring during normal brain development (BrainSpan database) using the *Singscore* method. We observe similar developmental patterns in LMNSC01 organoids and developing cortex for genes characterizing neurons, astrocytes, and oligodendrocytes, and multiple pathways including those involved in apoptosis, neuronal cytoskeleton, neurotransmission, and metabolism. Notable properties of this LMNSC01 platform are its initiation with immortalized authentic human neural stem cells, growth in a physiological oxygen environment, the consistency of the organoids produced, and favorable comparison of their gene expression patterns with those reported for normal cortical development.

**SUMMARY:** E*x vivo* studies of human brain development has been advanced by adoption of organoids recapitulating to varying extents *in utero* patterns of tissue development and cell differentiation. We here present an organoid-based human cortical development platform employing immortalized fetal neural stem cells (LMNSC01) grown in a physiological (4% oxygen) environment. Characterizing LMNSC01 organoids for over 120 days *in vitro* by immunofluorescence and expression profiling (using NanoString), and then comparing these profiles to those of normal cortical development (BrainSpan database), revealed similar developmental patterns for neurons, astrocytes and oligodendrocytes. Notable properties of this platform are its initiation with immortalized authentic human NSCs, growth at physiological oxygen concentration, and subsequent favorable comparison of their gene expression patterns with those observed during cortical development.

## INTRODUCTION

Brain organoids comprised of human cells recapitulating patterns of cell differentiation and morphological development characteristic of human brain ^1–3^ to a greater extent than monolayer or spheroid cultures are a powerful platform for *ex vivo* experimentation ^2,4,5^.

We here present a novel organoid platform based on L-*MYC*-immortalized fetal human neural stem cells: LMNSC01^6,7^. These cells can be expanded to create research cell banks while maintaining a stable genetic background ^7^. This report is concerned with a cohort of organoids, initiated with LMNSC01 neural stem cells and grown at physiological (4%) oxygen concentrations, that survive for more than four months without signs of necrosis. We observe by immunofluorescence and confocal microscopy emergence of morphologically distinct cells of multiple neural lineages (neuron, astrocyte, oligodendrocyte) expressing differentiation markers indicating cortical/telencephalon identity. At the same time, populations of SOX2-expressing neural progenitor cells are maintained with no evidence of uncontrolled proliferation.

We validated this LMNSC01 organoid platform by profiling gene expression in developing organoids (by NanoString), and determining quantitatively (by *Singscore* method) that these patterns compare favorably to those described for developing human brain (BrainSpan database). We also demonstrated organoid-to-organoid reproducibility by observing correlated expression of gene panels in individual organoids of the same age, and separation from those of different ages.

Developing platforms for *ex vivo* studies of human brain development and formulation of therapeutic strategies for brain disorders has been hampered by limited access to human brain tissue, inadequate *in vitro* models of developing human brain, and use of rodent models that imperfectly recapitulate human brain physiology ^8,9^. We suggest that LMNSC01 organoids provide a physiologically relevant and consistent platform for examining regulation of cortical development and function.

## RESULTS

### LMNSC01 organoids exhibit consistent growth and gene expression patterns

Differentiation of LMNSC01 cells into multiple types of neural lineage cells (i.e., neurons, astrocytes and oligodendrocytes) was originally described in monolayer culture^6^, and, here, in organoid cultures. Matrigel-imbedded LMNSC01 cells are grown as free-floating organoids in a 4% O_2_ environment approximating that present in fetal brain using characterized NeuroCult Proliferation medium. Over a 1- to 3-month period, LMNSC01 cells within the organoids, differentiate into cells expressing markers of maturing and mature neurons including cortical and hippocampal fated cells, radial glia, astrocytes, and oligodendrocytes, as well as endothelial cells. These organoids are most properly termed “guided” in that they are grown in a medium favoring proliferation (but not otherwise directive), and “telencephalic” based on their tissue of origin^10^.

In free-floating LMNSC01 organoids, as illustrated in the brightfield images in **Figure 1A**, **t**he neural stem cells initially proliferate to fill the space delimited by the Matrigel droplet (D0, D15), and then continue to expand, developing internal structure and in some cases distorting the spherical structure of the early organoids (D33, D47). This progression of organoid internal structure is also illustrated in **Figure 1B** using eGFP-expressing LMNSC01 cells. At D15 the NSCs appear distributed relatively homogeneously throughout the organoid, with a more complex internal structure evident at D30. An increasingly complex organization is evident at D60 and is maintained at D90, a self-organization promoted by the supporting Matrigel extracellular matrix.

**Figure 1.**
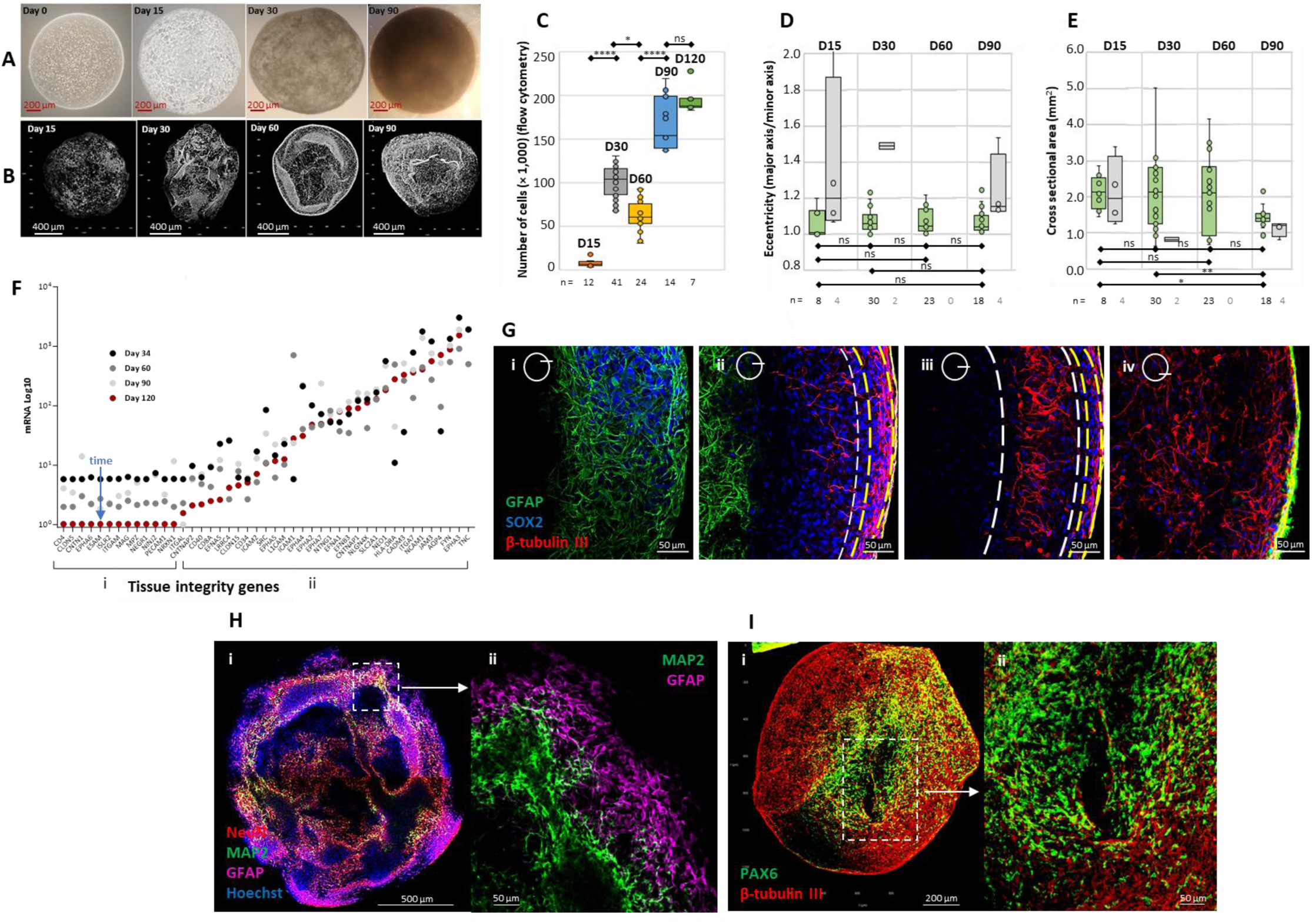
LMNSC01 organoid growth, maturation, and layer formation. **(A)** Brightfield images of LMNSC01 organoids from an initial seed of 1000 cells/organoid (D0). Organoids progressively increase in confluency between D0 and D47. At D15 the organoids appear more homogeneous; internal structure becomes evident at D30. **(B)** Reconstructed confocal images of enhanced green fluorescent protein (eGFP)-expressing LMNSC01 cells illustrating expansion, folding, and internal architectural complexity that becomes evident at D30. Some organoids curl (D60), while most remain roughly spherical (D90). **(C)** Increase in numbers of cells per organoid over 120 days in culture: rapid growth over the first 30 days, a characteristic dip at D60, followed by stabilization at D90 to D120 (*: p < 0.05; **: p < 0.01; ****: p < 0.001). Organoids were dissociated, and cells counted by flow cytometry. Cell viability after organoid dissociation: D15, 96.6 ± 1.0%; D30, 92.0 ± 2.6%; D60, 88.2 ± 2.4%; D90, 95.2 ± 0.3%; D120, 91.4 ± 0.5% (mean ± s.d., number of organoids as indicated on the figure). **(D)** Eccentricity of organoid shape assessed as major axis/minor axis, illustrating that most organoids were smooth and ovoid (green bars). Grey bars denote organoids identified by eye as not smoothly spherical or ovoid. By this measure, 1.0 indicates a spherical organoid. Numbers of organoids are as indicated on the figure. The majority of organoids were smooth spherical/ovoid: D15, 66%; D30, 94%, D60, 100%, D90, 82%. Statistical tests are between smooth/ovoid populations. **(E)** Organoid cross-sectional areas over time in culture. This value remained relatively stable at around 2 mm^2^ for both spherical/ovoid and more eccentric organoids. Mean diameters measured as average of major and minor axis (mm): D15, 1.7 ± 0.2; D30, 1.6 ± 0.4; D60, 1.6 ± 0.4; D90, 1.3 ± 0.1 (mean ± s.d.); p < 0.05 for all days relative to D90, all other comparisons not significant; numbers of organoids as indicated on the figure. **(F)** Expression (NanoString) of genes in the pathway “Tissue Integrity and Maintenance” (**Supplemental Table 1**), ordered by expression level at D120. **(i)** Genes that show their highest expression at D30, and whose expression becomes minimal at D120 (blue time arrow). **(ii)** Genes with higher expression in organoids between D34-D120 and whose expression is maintained at D120. **(G)** D30 organoid. Series of images of organoid edge at increasing depth (schematic), as described in the text, showing stratification of GFAP^POS^, SOX2^POS^, and β-tubulin III^POS^ cells. **(H)** D30 organoid. Neurons (NeuN^POS^ and MAP2^POS^) are internal to GFAP^POS^ glia near the organoid surface. **(I)** D30 organoid. PAX6^POS^ cortically fated cells surrounding a ventricle-like cell-free area, internal to a surround of β-tubulin III^POS^ immature neurons.

Over the first 120 days in culture, numbers of cells per organoid show rapid increase over the first 30 days, exhibit a characteristic dip at D60, and then stabilize at approximately 200,000 cells per organoid by D90-D120 (**Figure 1C**). Ultimately, as illustrated by measures of eccentricity (**Figure 1D**), the majority of organoids (greater than 82% for D30-D90) were smoothly spherical or ovoid, with D15 organoids being the most irregular in form (66% smooth/ovoid). The more spherical or ovoid D15-D60 organoids had relatively stable cross-sectional areas of around 2 mm^2^ (**Figure 1E**), with D15-D60 organoids having diameters of 1.6-1.7 mm and D90 organoids being somewhat smaller (1.3 mm). In this organoid system, size and thus differentiation may be constrained by Matrigel drop size ^11^ rather than oxygen and nutrient exchange. In larger organoids it is thought that cells further than 200-400 µm from the surface do not receive sufficient nutrients through diffusion ^12^ and healthy tissue can be limited to the organoid surface ^2^. However, LMNSC01 organoids survived for more than four months without signs of necrosis.

Interrogating expression of genes in the NanoString Tissue Integrity theme (**Supplementary Table S1**) over time (**Figure 1F**), and then ordering them according to expression level on D120, revealed two temporal patterns. In **group i**, expression of these genes was similar at D34 and ceased by D120, while expression of genes in **group ii** started at higher levels than those in **group i**, were consistently elevated at D120, and increased or decreased over the interval D34-D120. Gene ontology (GO) analysis (**Supplementary Figure S1)** suggests that there was an increase in representation of processes involved in shaping organoid architecture: “localization”, “interspecies interaction between organisms”, “developmental processes”, “locomotion”, “multicellular organismal processes”.

By D30, LMNSC01 organoids formed layers reminiscent of developing human cortex ^13^, as illustrated in a series of images at increasing depths along an organoid edge (**Figure 1G**). Towards the organoid interior (**panels i, ii**), GFAP-expressing cells mark a subplate-intermediate zone-like area containing radially aligned β-tubulin III neurons (**panels ii, iii**) and a cortical plate-like region (between white lines in **panel iii**). Towards the organoid surface, yellow lines demarcate a marginal zone-like region containing circumferential GFAP-expressing glia (**panels i, iv**) and β-tubulin III-expressing neurons (**panels ii, iii**) extending to the organoid surface (**panel iv**). This marginal zone-like pattern of NeuN- and MAP2-expressing neurons internal to GFAP-expressing glial cells is also illustrated in **Figure 1H**. In **Figure 1I**, a subventricular zone-like area of area of PAX6-expressing cells surrounds a cell-free ventricle-like structure.

### Cells in LMNSC01 organoids express markers of cortical and hippocampal neural identity

As shown in **Figure 2**, LMNSC01 organoids expressed markers indicative of a telencephalon (cortical and hippocampal) identity. SOX2-expressing stem and progenitor cells were distributed throughout the organoids, decreasing as a percentage of total cells between D15 and D60, and stabilizing at about 80% of cells by D120 (**Figure 2A1, A2**). FoxG1, marking forebrain-fated cortical progenitor cells ^14^ were found in D30 organoids and less so in organoids at D90 (**Figure 2B**). Wnt receptor Fzd9, marking hippocampal fate ^15^, was strongly expressed throughout organoid development, even as cell shapes matured (**Figure 2C**). Finally, transthyretin (TTR), marking choroid plexus fate ^16^, was also found in D30 organoids (**Figure 2D**). NanoString expression profiling of organoid cell type markers (**Figure 2E**), and consolidated into a dendrogram (**Figure 2F**), indicates the closest relatedness of organoids to cortical tissue, and their distance from other regions.

**Figure 2.**
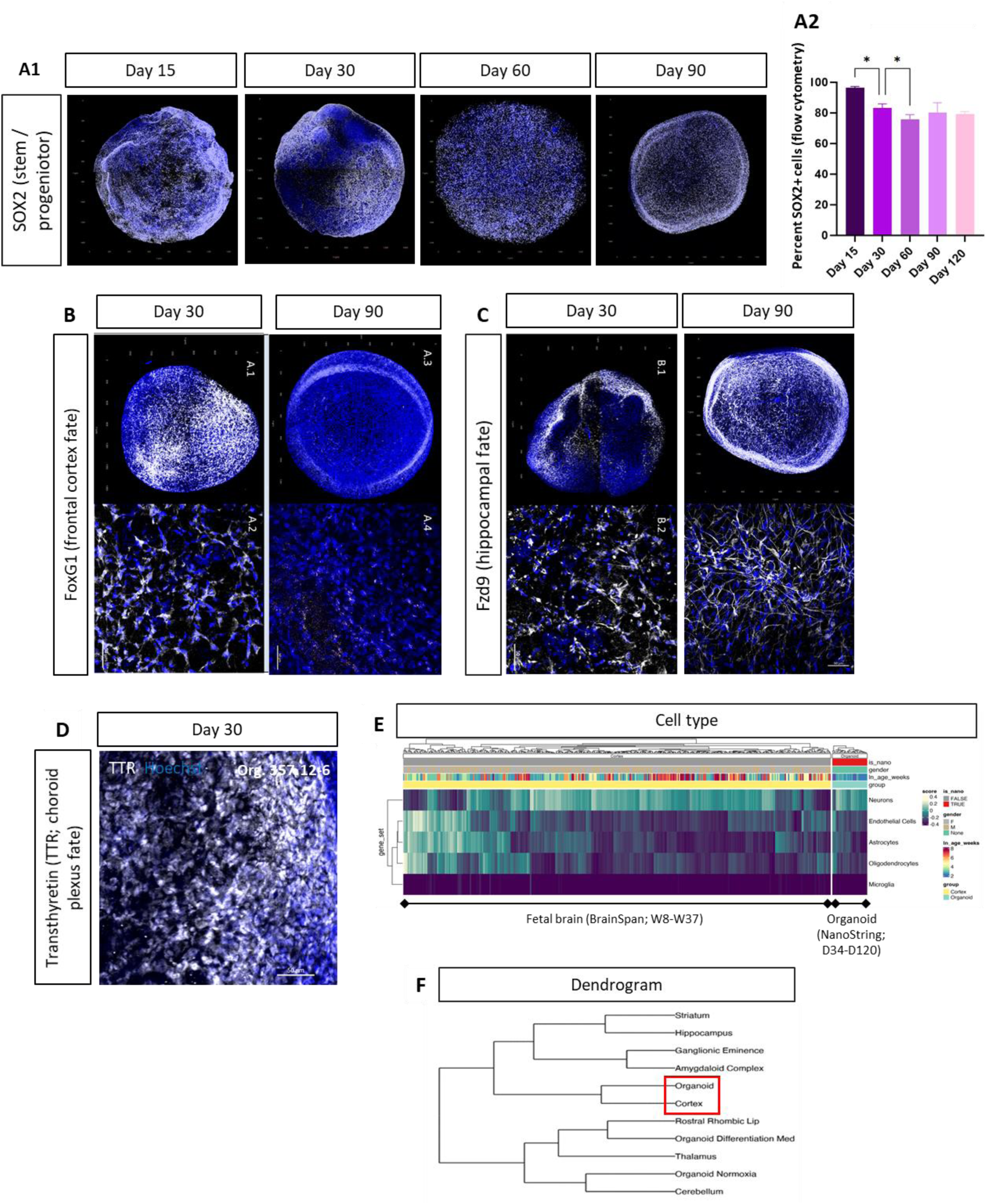
LM-NSC008 organoids express markers indicative of a telencephalon (cortical and hippocampal) identity. **(A1)** 3D reconstruction of organoid immunofluorescence; SOX2-expressing stem and progenitor cells remain distributed throughout organoids over the period D15-D90. **(A2)** Percentage of SOX2^POS^ cells over the *in vitro* period (D15-D120), as measured by organoid dissociation and flow cytometry. *: p < 0.05, n = 4 at each time point. **(B,C)** Paired images of entire organoids (3D reconstruction, above) and detail (single optical section, below). **(B)** FoxG1 immunofluorescence marking forebrain-fated cortical progenitor cells. **(C)** Persistence of Wnt receptor Fdz9, a marker of hippocampal fate, D30-D90, also illustrating changes in cell morphology. **(D)** Expression of TTR (a marker of choroid plexus fate) by immunofluorescence. D30 organoid. **(E)** Heatmap of cell type clustering by expression profiling within fetal cortex (BrainSpan, in weeks post-conception) and organoids (NanoString, in weeks *in vitro*). **(F)** Dendrogram constructed from NanoString expression profiling of cell type markers (from **E**), indicating the closest relatedness of organoids to cortex and distance from other regions.

### Cells in LMNSC01 organoids differentiate into multiple types of neural lineage cells

Differentiation of LMNSC01 cells into neurons, astrocytes and oligodendrocytes was originally described in monolayer culture ^6^. LMNSC01 cells grown in 3D organoid culture showed robust differentiation into NSC lineages as presented in roughly developmental order in **Figure 3** by immunofluorescence (**Figure 3B1-E1**), as percentages of total cells (by dissociation and flow cytometry, **Figure 3B2-E2**), by organoid mRNA expression (NanoString, **Figure 3A2-E2**), and comparison to fetal cortical mRNA (BrainSpan, **Figure 3A2-E2**).

**Figure 3.**
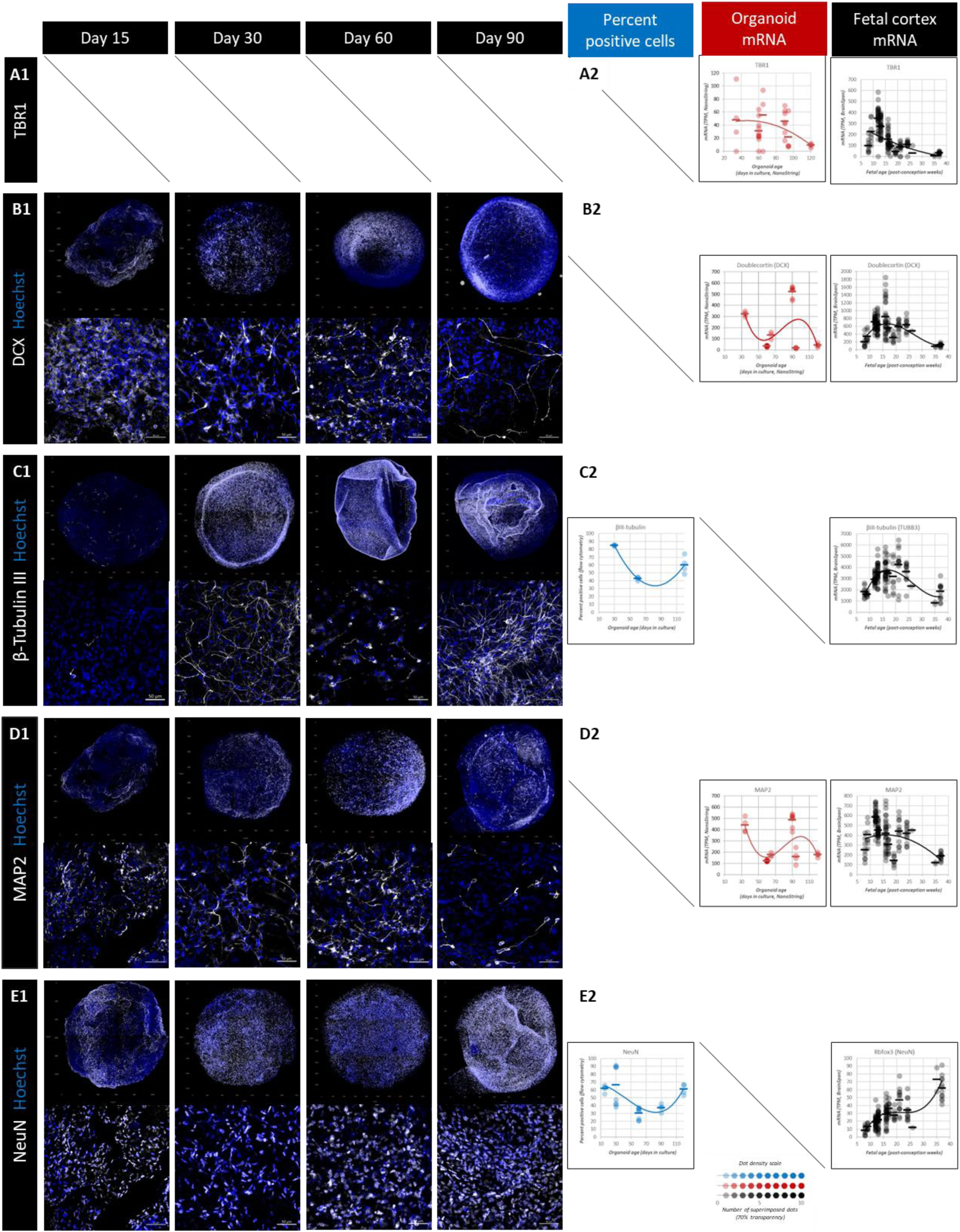
Progression of neuronal differentiation marker expression over time. *Diagonal lines indicate image or data not available*. **(A1-E1)** Immunofluorescence image pairs of representative organoids. Above, confocal 3D reconstructions of entire organoids; below, detailed higher resolution single optical sections (below). **(A2-E2)** Expression of markers, by percentage of total positive cells (by dissociation and flow cytometry; blue), and by mRNA levels in organoids (NanoString; red) and fetal cortex (BrainSpan; black). Data are plotted at 70% transparency; the inset illustrates increasing density of dots with overlap of multiple points. **(A2) Transcription factor T-box brain1 (Tbr1)**, a marker of postmitotic neurons ^17^, see text. **(B1, B2) Doublecortin (DCX)**, a marker of migrating early neurons, see text. **(C1, C2) β-Tubulin III**, a marker of maturing neurons ^19^; see text. **(D1, D2) MAP2**, a neural differentiation marker ^20^; see text **(E1, E2) NeuN**, a nuclear marker of mature neurons ^21^; see text.

Expression of transcription factor T-box brain1 (Tbr1) mRNA, marking postmitotic neurons ^17^ (**Figure 3A2**), was higher at early times in organoids (NanoString) (D34-D60/65), and fell progressively (D94-D120) as organoids aged. In fetal cortex (BrainSpan), expression was low at W8/9, elevated at W12/13, and then fell over the interval W16/17 to W35/37. Immunofluorescence also indicated Gremlin 1, important for early cortical excitatory neuron development ^18^, was expressed in the LMNSC01 organoids (not shown).

Doublecortin (DCX), a marker of migrating immature neurons, was present throughout organoid development, with, by immunofluorescence, relative numbers of DCX-expressing cells appearing to decrease between D15 and D90 (**Figure 3B1**). At the mRNA level, expression in organoids (NanoString) was elevated at D34, transiently dipped at D60/65, was high at D90, and then declined over the interval D94 to D120. In developing cortex (BrainSpan), expression increased over the interval W8 to W16/17, then declined over the interval W19 to W35/37 (**Figure 3B2**).

Expression of β-tubulin III, marking maturing neurons ^19^ appeared in organoid immunofluorescence images to increase between D15 and D30, decrease somewhat in the D30-D60 interval, and again increase by D120 (**Figure 3C1).** Counts of percent positive cells **(Figure 3C2)** followed a similar pattern, elevated at D30, reduced at D60, and again elevated at D120. The morphology of β-tubulin III-expressing cells varied over time, becoming longer and more extended over the period D30 to D90, (**Figure 3C1**), perhaps reflecting tubulin’s role as a cytoskeletal protein. In fetal cortex (BrainSpan), β-tubulin III expression, similar to DCX, increased somewhat from W8 to W16-W17, and then decreased from W24/26 to W35/37 **(Figure 3C2).**

MAP2, a neural differentiation marker ^20^, was also present from the earliest times (**Figure 3D1**). At the mRNA level, (**Figure 3D2**) expression in organoids (NanoString) was biphasic, elevated at D34, reduced at D60/65, transiently increased at D90, and reduced at D94-D120. In developing cortex (BrainSpan) MAP2 expression (BrainSpan) was mixed during W8 to W16/17, and then progressively decreased from W24/26 to W35/37 (**Figure 3D2**).

Expression NeuN, a nuclear marker of mature neurons ^21^, was surprisingly high by immunofluorescence at D15 (**Figure 3E1**) and as a percent of total cells (**Figure 3E2**), appearing to decrease between D30 and D60, recovering by D90, and increasing to D120 (**Figure 3E2**). Fetal cortex NeuN mRNA levels (BrainSpan) increased from W8 to W16, paused from W17 to W26, and then increased until W35/37, ending the fetal period at higher levels (**Figure 3E2**).

There were some consistent expression patterns of these neural antigens. In organoids, markers reflecting intermediate stages of neural differentiation (DCX, β-tubulin III, MAP2) all showed a dip [by cell counts and mRNA (by NanoString)] centered on D60, which is also the time of slower organoid growth as illustrated in **Figure 1C**, before recovering around D90 and, for the mRNA measurements (NanoString), decreasing again at D120. In fetal cortex (BrainSpan), expression of these same three markers described an inverted-U, lower at W8/9, a local maximum at W16 to W20, and then reduction by W35/37. The earliest and latest developmental markers, Tbr1 and NeuN, exhibited reciprocal behaviors. Tbr1 ^17^ was expressed in organoids (NanoString) initially elevated at D30-D60 and lower at D120. Similarly, in fetal cortex (BrainSpan), Tbr1 expression was higher at W16 and lower by W35-W37. NeuN, marking later stages of neural maturation ^21^, was in organoids (by cell counts) initially high at D15 (a possible indication of precocious maturation), showed the characteristic dip at D60-D90, before increasing by D120. In fetal cortex NeuN expression (BrainSpan) was reduced at W8/9, increased to W16, stabilized between W17-W26, and then was elevated by W35/37.

LMNSC01 organoids also incorporate macroglia: astrocytes and oligodendrocytes. As illustrated in **Figure 4**, SOX9, a transcription factor expressed by neural progenitor cells and by astrocytes in adult brain ^22^, is found in D30-D60 organoids by immunofluorescence, most notably in the nuclei of vimentin-expressing cells (**Figure 4A, Dai-iii**). For oligodendrocytes, immunofluorescence shows expression of maturing oligodendrocyte marker O1 from D30 to D60 in organoids **(Figure 4B)**. Phosphorylated vimentin, a marker of mitotic astroglial cells ^23^, was present in cell bodies, showed an increase in expression from D15 to D30 and then declined by D90 (**Figure 4C)**. The astrocytic cytoskeletal marker GFAP was present in organoids through the entire culture period to D120, with cells appearing more fibrous at later times (**Figure 4Daiv-vi)**

**Figure 4.**
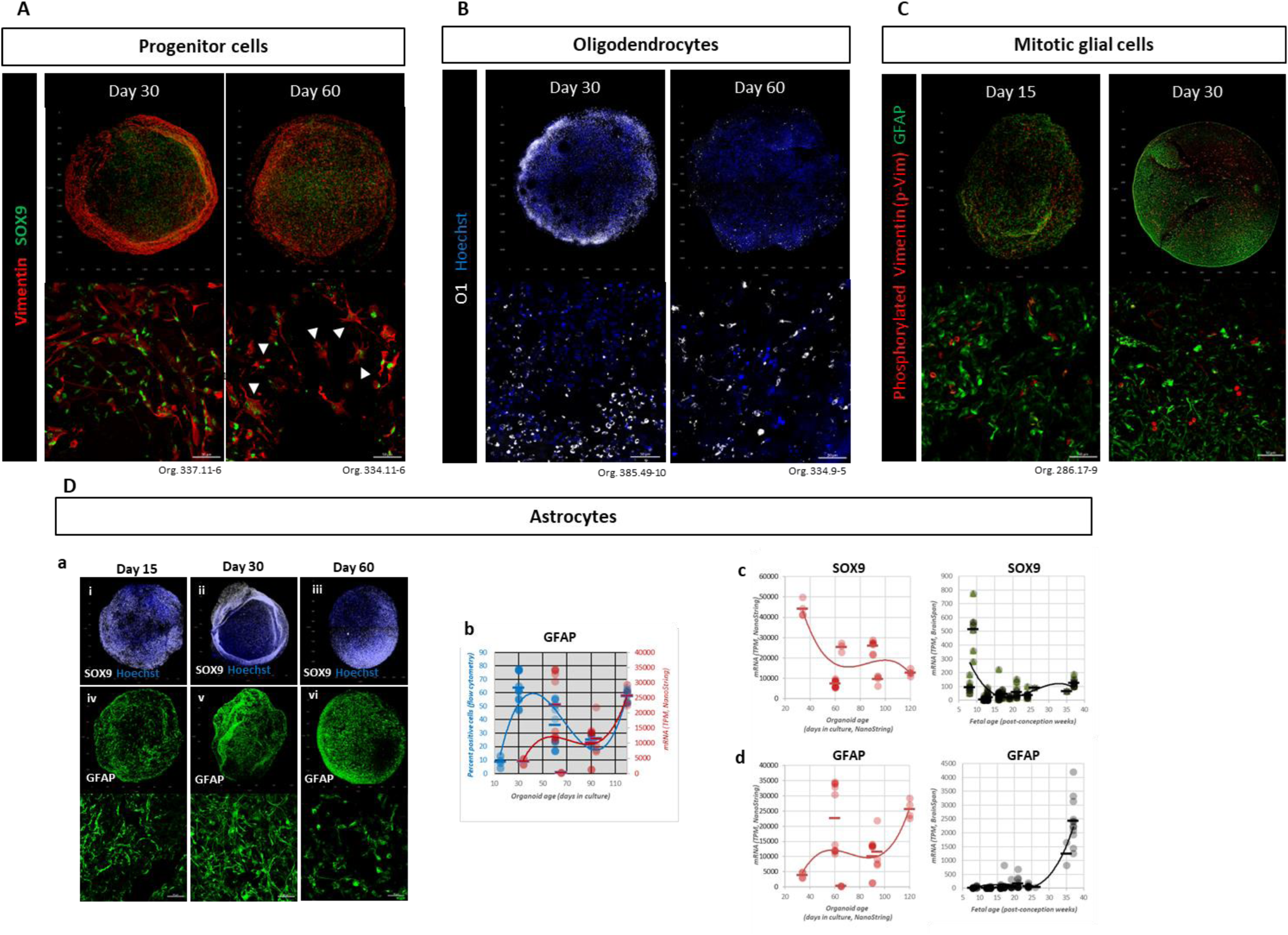
Differentiation of macroglia (astrocytes and oligodendrocytes) in organoids over time. *In this figure, each dot represents a data point from an organoid (NanoString) or a cortical specimen (BrainSpan). The horizontal lines indicate the mean of the data. The lines are included to aid in viewing trends in the data*. **(A)** Expression of SOX9 by immunofluorescence, often in the nuclei of vimentin-expressing neuroprogenitor cells (white arrow heads). **(B)** Expression of oligodendrocyte marker O1 by immunofluorescence. **[C]** Expression of phospho-vimentin in cell bodies of mitotic GFAP-expressing cells by immunofluorescence. **(Da, i-iii)** Maintained expression of SOX9 in organoids of increasing age by immunofluorescence. **(Da, iv-vi)** Expression of GFAP in organoids of increasing age by immunofluorescence. **(Db)** Almost parallel expression of GFAP as percent expressing cells (flow cytometry) and mRNA (NanoString), with a transient decrease around D60. **(Dc)** Expression of SOX9 mRNA in organoids (red, NanoString) and fetal cortex (black, BrainSpan). **(Dd)** Expression of GFAP mRNA in organoids (red, NanoString) and fetal cortex (black, BrainSpan). Numbers of organoids expression profiled by NanoString for days in culture (D); numbers of organoids/number of cohorts: D34, 4/1; D60, 10/2; D65, 4/1; D90, 7/2; D94, 4/1; D120, 4/1. Numbers of cortical region fetal samples as reported by BrainSpan for weeks post-conception (W); numbers of samples/number of donors: W8, 9/1; W9, 7/1; W12, 31/3; W13, 32/3; W16, 28/3; W17, 9/1; W19, 8/1; W21, 11/2; W24, 11/1; W25, 1/1; W26, 3/1; W35, 1/1; W37, 11/1.

Patterns of GFAP expression in organoids, assessed as percent GFAP^POS^ cells (by flow cytometry) and as mRNA levels (by NanoString) were essentially parallel (**Figure 4Db**), showing increase through D30 to D60, a dip at D90, and an increase at D120. We also observed reciprocal maturation patterns of expression of the progenitor marker SOX9 and the more mature marker GFAP in both organoids (by NanoString) and fetal cortex (by BrainSpan) (**Figure 4Dc**,**4Dd**). Reciprocal expression of genes characteristic of immature and mature cell states has been noted in other brain organoid studies. For example, Lancaster et al. (2013) observed neuronal differentiation (increase in DCX^POS^ cells) at the expense of SOX2^POS^ progenitors in microcephaly models ^2^.

### LMNSC01-derived organoids express mesodermal markers of endothelial cells and of microglia

As illustrated in **Figure 5**, we noted intrinsic expression of endothelial markers CD31 (PECAM-1) and von Willebrand Factor (vWF), as well as tight junction marker ZO-1, in organoids created with only LMNSC01 cells. This apparent appearance of mesodermal lineage cells was noted as early as D15. While expression of these markers was less organized at D15, in older organoids (D30, D90) they tended to be seen together and could be observed coalescing in linear arrangements reminiscent of nascent vasculature (yellow arrowheads in **Figure 5**). This immunochemistry was consistent with expression of the individual mRNAs comprising the NanoString endothelial pathway throughout organoid development, declining somewhat in older D120 organoids as well as in the aggregate *Singscore* **(Supplemental Figure S2**). Expression of these same genes was also evident in developing cortex (BrainSpan) (**Supplemental Figure S2**), the increases at later times possibly relating to endothelial cell invasion during cortical vascularization ^24^.

**Figure 5.**
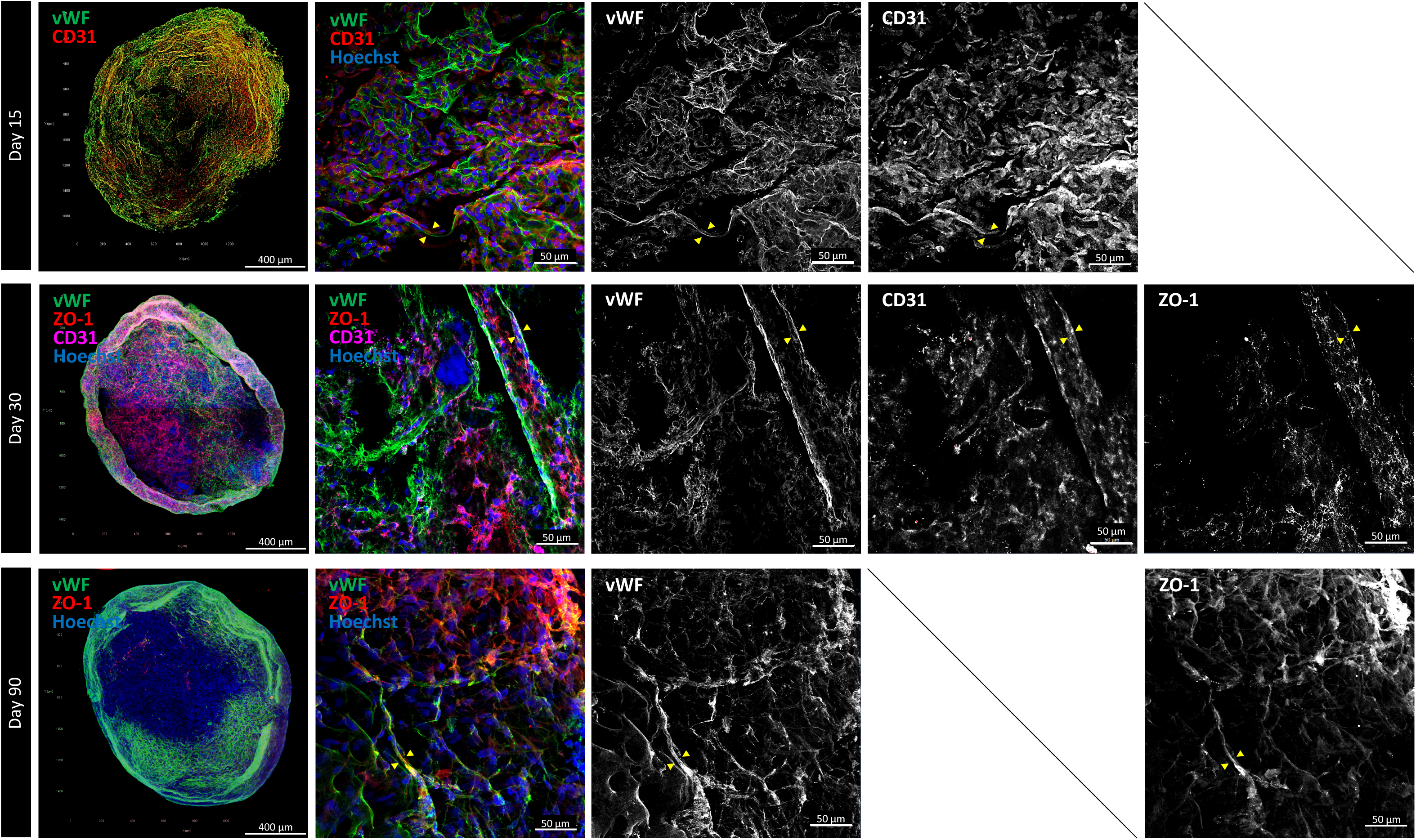
Immunostained LMNSC01 organoids express endothelial markers CD31 and von Willebrand Factor (vWF), and the tight junction protein ZO-1. Yellow arrowheads indicate alignment of these markers in linear arrangements suggestive of nascent vasculature, increasing in older organoids.

Emergence of vascular-like structures may not be unexpected. Human vasculature is formed using soluble factor- and contact-dependent interactions between neural stem cells and endothelial cells ^25^, conditions that could arise in the organoids. Additionally, apparent direct transdifferentiation of neural stem cells to endothelial-like cells has been reported ^26^. Gliobastoma cells also directly undergo apparent endothelial transdifferentiation *in vitro* and in *in vivo* xengraft models ^27^.

mRNA markers of microglia were also present in LMNSC01 organoids (by NanoString) (**Supplemental Figure S2**), an unexpected observation given their mesodermal origin in yolk sac erythromyeloid progenitors ^28^. The low abundance of microglia in developing cortex (0.5-1% of cells compared to 10% of cells in mature brain) appears reflected in the low mRNA expression of marker genes in developing cortex (BrainSpan) as compared to the organoids (NanoString) (**Supplemental Figure S2**). Similar intrinsic differentiation of microglia in cortical organoids was observed by Ormel et al ^29^, as was appearance of mesodermal progenitors ^30^.

### Temporal progression of developmental programs in organoids compares favorably to fetal cortical development

As presented in **Figure 6**, we used bulk RNA expression profiling to examine developmental programs in organoids over time in culture ^31^. Shown in **Figure 6A1** is cross correlation for expression of the genes in the NanoString Neuroinflammation panel (**Supplemental Table S1**) by each of 33 organoids ranging from D34 to D120 in culture. Evident are consistent high correlations between organoids of the same ages, and sharp demarcations between organoids at other ages, an indication of stable *in vitro* development distinguished by time in culture. **Figure 6A2** shows a similar cross correlation for the same genes during normal fetal development (BrainSpan). As with the organoids, but not as dramatically, expression of the gene set in cortex was consistent with developmental timing.

**Figure 6.**
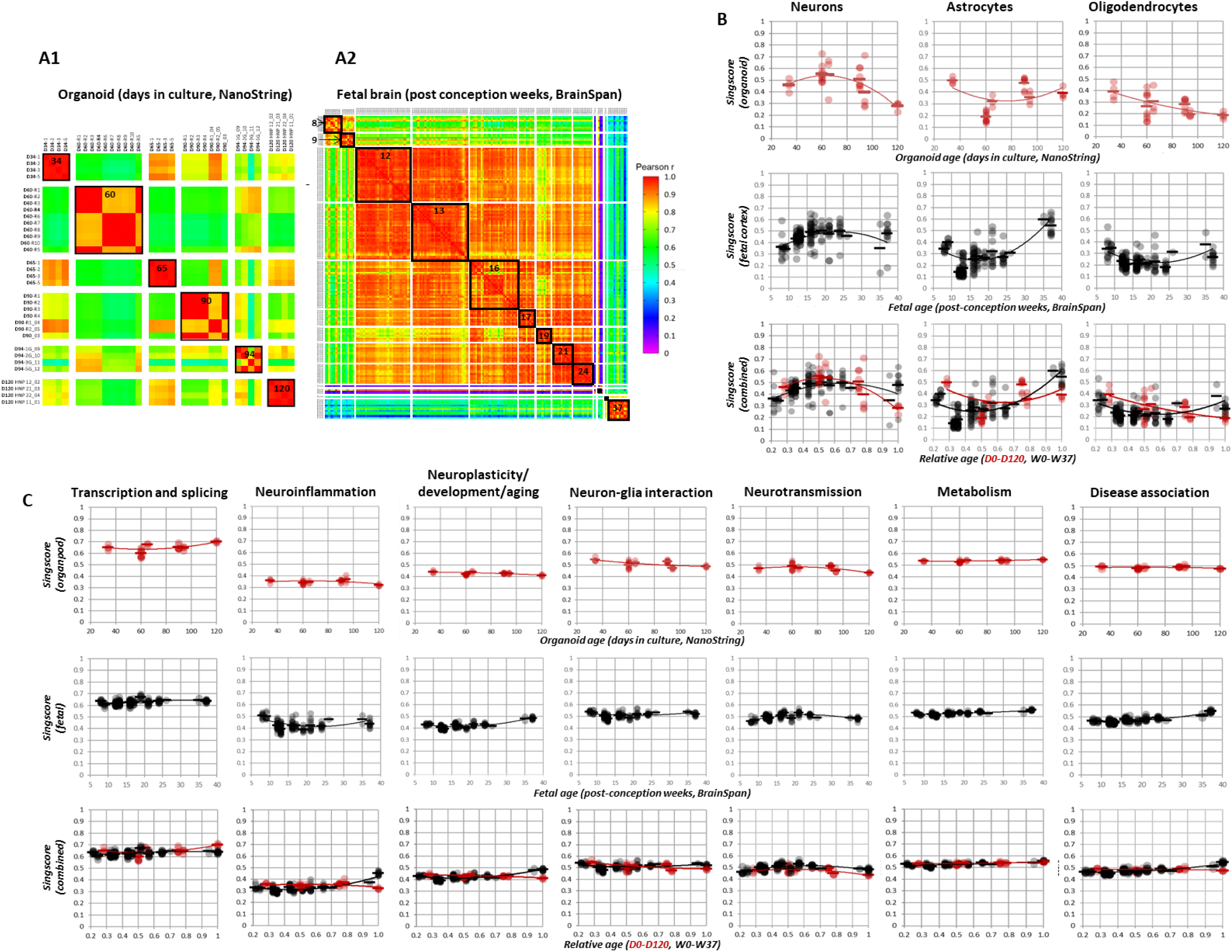
Progression of developmental programs in organoids and during fetal cortical development. **(A1)** Cross correlation for expression of the genes in the NanoString Human Neuropathology Panel (**Supplemental Table S1**) by each of 33 organoids ranging from D34 to D120 in culture, illustrating the similarities in gene expression between organoids of similar ages, and pronounced differences with organoids of different ages. **(A2)** Similar cross correlation for the same genes during normal fetal development drawn from the BrainSpan database. **(B)** *Singscores*, expression relative to NanoString-defined reference genes, where 0.5 indicates no difference, over time for three major NSC-lineage cell types in organoids (top, red) and fetal cortex (middle, black). Below are the same data plotted on a relative time scale, 0.0-1.0, for organoids (D0-D120 in culture) and fetal cortex (W0-W37 post-conception, W40 marks full human gestation). These plots illustrate the quantitative similarities between organoids and fetal cortex for both expression levels and waveforms. **(C)** Similar data for seven themes and pathways (**Supplementary Table S1**); the complete data are shown in **Supplemental Figure S2**.

As presented in **Supplemental Table S1,** NanoString organizes genes into six fundamental themes, 24 pathways, and five cell types. To facilitate direct comparison of *in vitro* and *in vivo* development, we normalized both sets of data to the NanoString internal reference genes (**Supplemental Table S1)**, with *Singscore* values centered on 0.5 indicating deviation from these reference genes as generated by the *Singscore* algorithm ^32^. *Singscore* values for several examples of cell types and developmental themes of particular interest are shown in **Figures 6B** and **6C**; the complete set of comparisons is presented in **Supplemental Figure S2**. Evident is that for neurons, astrocytes and oligodendrocytes, expression levels (relative to reference genes) and the velocity of changes for organoids between D34 and D120 (red), and fetal brain between W8 and W37 after conception (black), appear very similar. We examined this concept more explicitly by constructing proportional time scales between 0.0 and 1.0 in which the day of organoid initiation and the day of conception were time 0.0, and the oldest organoids we examined (120 days), and 37 weeks post conception (close to the 40 weeks of normal gestation in humans) were time 1.0. In this scheme, the data could be remarkably similar, as indicated in the overlays of these curves, suggesting that for many processes, development of LMNSC01 cells in organoids was paralleling the normal human progression in cortex.

## DISCUSSION

Here, we present an organoid platform of *in vitro* telencephalon development formed from human fetal neural stem cells (NSCs) immortalized by expression of L-*MYC.* This organoid platform differs from the most commonly used brain organoids in that they employ human NSCs grown in a 4% O_2_ environment that more faithfully reproduces the *in vivo* context of developing human cortex ^33^. In contrast, most brain organoids are grown in ambient room air with 20% O_2_ present, or, rarely, in high-oxygen environments (40%) ^34^. Growth in the elevated O_2_ concentration of room air may have consequences for organoid development, as the more physiological 4% O_2_, similar to O_2_ levels of 1-5% reported for developing brain ^33^, favors self-renewal and neurogenic potential through activation of Wnt signaling pathways ^33,35^. It is only the severe hypoxia associated with ischemia (less than 1% O_2_) that results in brain damage ^33^.

The LMNSC01 cells employed here are well-characterized for NSC lineage differentiation ^6,7^. Organoids derived using LMNSC01 cells displayed reproducible patterns of proliferation, identity, and associated structural organization, for over 100 days in culture, and survived for more than four months without signs of necrosis. By immunochemistry and bulk RNA expression profiling we were able to assess the cellular compositions of individual organoids ^36^, and demonstrate gene expression patterns unique to organoid age (**Figure 6A**), an indication of coordinated developmental progression over time. The consistent reproducibility of these organoids is an important factor in achieving the full utility of this platform ^37–39^.

Transcriptome analysis by NanoString showed time-dependent changes in RNA expression patterns for genes characterizing neurons, astrocytes, and oligodendrocytes, as well as generalized developmental processes, that compared favorably, both quantitatively and temporally, to those seen in human cortex during fetal development, especially at early times (**Figure 6B**, **6C**, **Supplemental Figure S2**). Camp et al ^40^ also noted similarities between gene expression patterns of organoids and normal fetal development, reflecting a consensus that iPSC-derived organoids reflect cortical development as we concluded here for the LMNSC01 organoids.

Nevertheless, establishing human brain organoids directly from NSCs obviates some of the challenges that have emerged when employing embryonic stem cells (ESCs) or induced pluripotent stem cells (iPSCs) as starting materials. These include, depending on the protocol and on the desired neural cell type, potentially months to achieve iPSC reprogramming ^41,42^, as well as incomplete or erroneous reprogramming ^43^, possibly related to cell of origin ^44^, resulting in defects in DNA methylation, chromatin regulation, protein synthesis ^45^, poor or incomplete differentiation ^4,45–49^, as well as accumulating somatic mutations ^50^ resulting in organoid heterogeneity ^51,52^ that compromise their ultimate utility ^37^.

The human NSC line utilized here (LMNSC01) reflects the normal phenotype of NSCs at their time of immortalization (12 weeks of gestation)^6^, rather than of adult NSCs as seen for granule cells of the hippocampal dentate gyrus or the progenitor cells in the rostral migratory stream. As such, they may not appear as versatile as the induced pluripotent stem cells (iPSCs) that have emerged as a source of neural stem or progenitor cells (NSCs or NPCs) used to establish brain organoid cultures using cells from patients with inborn genetic neurological and other conditions ^42,53,54^. However, preliminary studies from others have demonstrated that gene editing techniques can introduce precise changes in genetic makeup of NSCs underlying these disorders ^55–57^. Taken together with their stability, these studies suggest LMNSC01 organoids as an alternative approach to establishing models of inborn brain diseases.

The consistency and reproducibility of LMNSC01 organoids, together with their recapitulation of fetal expression patterns, provide the opportunity to observe and control the development and differentiation of multiple neural-lineage cell types within a controlled culture system, assess the impact of environmental perturbations on these processes, as well as serve as a platform for drug and therapy screening. Further, their chromosomal stability and the absence of tumorigenicity make these organoids potentially suitable source material for implantation and engraftment therapies.

Responses of LMNSC01 organoid to variations in growth conditions further suggest future refinements that more closely following fetal brain development, as seen with, for example, the transition from NC Proliferation medium to NC Differentiation medium after initial periods in culture, as illustrated in **Figure 7** (see also ^2^), where changes in cell development after transition to a differentiating medium at D15 are illustrated (**Figure 7A**). Under this condition, organoids undergo growth arrest without a concomitant reduction in size (**Figure 7B**). By D30, β-tubulin III-expressing neurons (**Figure 7C)** develop varicosities in differentiating medium (**i → ii**) suggesting synaptogenesis, as indicated by expression of synapsin1 in MAP2-expressing neuron processes (**iii**). By D104, GFAP^POS^ astrocytic cells (**Figure 7D**) in differentiating medium extend processes and form fascicles not evident when grown in NC Proliferation medium. O1-expressing oligodendrocyte-like cells (**Figure 7E**) in differentiating medium between D30-D104 acquire a ramified morphology also not evident when grown in proliferation medium (compare with **Figure 4B**).

**Figure 7.**
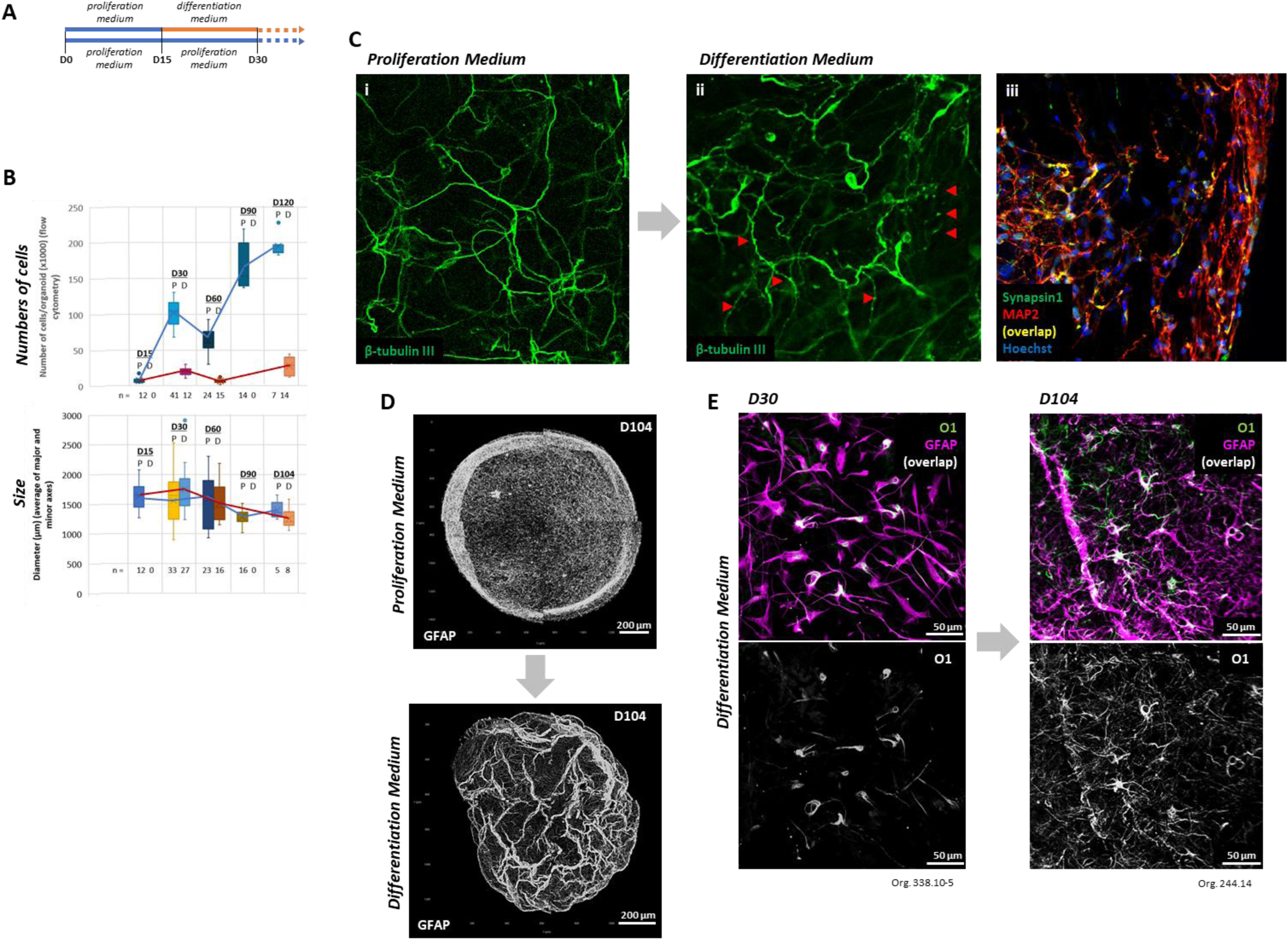
Growth conditions alter LMNSC01 organoid differentiation. **A,** Experimental schema. Organoids were initially grown (4% O_2_) in NeuroCult (NC) Proliferation medium until D15, then optionally transferred to NC Differentiation medium until D30 or longer. P = NC Proliferation medium. D = NC Differentiation medium. *n* = numbers of organoids as indicated. **B,** Numbers of cells per organoid (*above*) after D15 continued to increase in NC Proliferation medium (EGF and FGF supplemented) (blue line), but stabilized in NC Differentiation medium (no added EGF/FGF) (red line). Organoid diameters (*below*), averages of major and minor axes, were relatively independent of growth medium. **C,** Maturation of neuron morphology and apparent synaptogenesis at D30 in NC Differentiation medium. **i,** Neuronal processes, marked by β-tubulin III expression, were extended and smooth in NC Proliferation medium. **ii**, Neuronal varicosities developed in NC Differentiation medium. **iii,** Varicosities (MAP2^POS^) also express synapsin1. **D,** At D104, GFAP^POS^ astrocytes extended processes and formed bundles in NC Differentiation medium. **E,** Organoids grown in NC Differentiation medium from D15: O1^POS^ oligodendrocyte cells extended process by D30, and by D104 became highly ramified.

Additional refinements could add additional cell types providing a closer relatedness to fetal cortical development. For example, rather than growing progenitors and immature neurons on an initially homogeneous scaffold, introducing structural, cues or providing local presence of additional cell types may shape more physiological spatial arrangements of differentiated cells. Further, apparent *de novo* appearance of endothelial cells and vasculature-like structures, as well as microglial gene expression suggesting development of these cells, raise the possibility that culture conditions could be manipulated to promote appearance of these cell types, thereby providing a more complete cortical model and further enhancing the utility of these organoids ^58–60^.

## EXPERIMENTAL PROCEDURES

### Resource availability

Further information and requests for resources and reagents should be directed to and will be fulfilled by the lead contact, Michael Barish.

### Expansion of LMNSC01 cells

Fetal brain neural stem cells were obtained from discarded tissue from Advanced Bioscience Resources under a City of Hope Institutional Review Board approved protocol (City of Hope IRB 10079) and L-*MYC* immortalized as described previously ^6^. LMNSC01 cells were expanded using a hollow fiber bioreactor (Terumo Quantum Cell Expansion System) in serum-free NSC medium (RHB-A medium; Cell Science) supplemented with growth factors (10 ng/mL bFGF, 10 ng/mL EGF; both from PeproTech), 2 mM L-glutamine (Invitrogen), Gem21 NeuroPlex Serum-Free Supplement (GeminiBio-Products 400-160) and penicillin-streptomycin (Mediatech, 30-002-CI) ^61^.

### Organoid preparation

Prior to organoid assembly, expanded LMNSC01 cells were cultured to 80-90% confluence beginning at 10,000 cells/cm^2^ using NeuroCult Proliferation medium (StemCell Technologies 05751) supplemented with EGF (10 ng/mL, PeproTech AF-100-15), FGF (10 ng/mL, PeproTech 100-188), heparin (1 U/mL, (StemCell Technologies 07980), at 37 °C in a 4% O_2_, 5% CO_2_ environment. This medium was also used to culture the organoids.

Our method for LMNSC01 organoid preparation is modified from those described by Hubert et al ^62^ and Lancaster et al ^63^. Each well of a 6-well plate was prepared by placing a 2.5 × 2.5 cm square of parafilm, sterilizing with 70% ethanol, and placing it uncovered in a culture hood under UV light for approximately 30 min for the ethanol to evaporate. At the same time, 500 µL of growth factor reduced Matrigel (Corning 356231) was placed in a microcentrifuge tube to which 83,000 cells were added and gently mixed. Using a repeating pipettor (Eppendorf 4049-7134), ten 6 µL-droplets/well, each containing 1,000 LMNSC01 cells, were dispensed onto the parafilm, and allowed to polymerize for 3 hours in the 37 °C, 4% O_2_, 5% CO_2_ incubator. The parafilm inserts were then gently removed, and 4 mL of NeuroCult Proliferation medium supplemented as above was added to each well, into which six LMNSC01/Matrigel droplets were added using a sterile spatula. The culture plate was returned to the 37 °C, 4% O_2_, 5% CO_2_ incubator for three days, after which the culture plate was transferred to a sterilized orbital shaker maintained in the incubator at 100 rpm. Culture medium was changed once per week for the first three weeks, and then twice per week thereafter. In some cases, LMNSC01 cells were engineered to express lentiviral eGFP and used to create organoids for imaging 3-dimensional organization (Zen 3.6, Zeiss)

### Immunostaining and imaging

Organoids were fixed for 30 minutes in 4% paraformaldehyde (PFA) in PBS at room temperature (RT), and then washed three times (3×) for 15 min with Tris-buffered saline (TBS) containing 0.1% Triton X-100 (Tx-100) (TBS wash solution). After washing, samples were permeabilized and blocked for one hour in blocking solution: TBS containing 1% Triton and 5% high avidity bovine serum albumin (BSA). Organoids were then incubated overnight at 4 °C with primary antibodies diluted in the same TBS blocking solution, then washed 3× (15 min each) with TBS wash solution with gentle agitation. Secondary antibodies were then applied in TBS blocking solution and incubated overnight at 4 °C. Organoids were then washed 3× with TBS wash solution and mounted on a glass slide with Prolong Glass (Invitrogen). After at least 48 hr at room temperature in the dark to allow for curing of the Prolong Glass, organoids were imaged using Zeiss LSM-700 or LSM-900 confocal microscopes. Images were processed and analyzed using Zen 3.6 (Zeiss) and Imaris 9 (Oxford Instruments).

### Reagents for immunostaining

Primary antibodies used for immunofluorescence: Stem cells: SOX2 (EMD Millipore AB5603, 1:300). Astroglial lineage: GFAP (Abcam ab53554, 1:300); vimentin (Abcam ab8069, 1:200); phospho-vimentin (p-vimentin, Abcam ab22651, 1:700); SOX9 (Novus Biologicals AF3075, 1:100). Oligodendroglial lineage: O1 (ThermoFisher 14-6506-82, 1:300). Neural lineage: Gremlin 1 (Bioss Antibodies bs1475R, 1:100); doublecortin (DCX, Santa Cruz Biotech SC-8066, 1:50); β-tubulin III (Biolegend Tuj1 801202, 1:700); NeuN (Abcam ab104224, 1:150); MAP2 (Abcam ab11267, 1:300). Hippocampal progenitor: Fzd9 (Origene TA340718, 1:300). Cortical progenitor: PAX6 (Biolegend 901301, 1:300); FoxG1 (Abcam ab18259, 1:300). Endothelial differentiation: CD31 (PECAM1, Santa Cruz Biotech SC1506, 1:300); von Willebrand Factor (vWF, Dako A0082, 1:300); ZO-1 (Invitrogen 33-9100, 1:300). Choroid plexus: transthyretin (TTR, Serotec AHP1837, 1:100)..

Secondary antibodies: Donkey anti-rabbit Alexa Fluor 488 (Invitrogen A21206, 1:1000). Donkey anti-mouse Alexa Fluor 555 (Invitrogen A21147, 1:1000); Donkey anti-sheep Alexa Fluor 555 (Invitrogen A21436, 1:1000). Donkey anti-goat Alexa Fluor 647 (Invitrogen A21447, 1:1000). Donkey anti-mouse Alexa Fluor 647 (Invitrogen A31571, 1:1000).

Nuclear stain: Hoechst 33342 (Invitrogen H3570, 1:1000).

### Flow cytometry

Organoids were dissociated using Neuron Dissociation Solutions (Fujifilm Wako Pure Chemical 291-78001), consisting of Enzyme Solution (295-78021), Dispersion Solution (298-78011) and Isolation Solution (202-78031), per the manufacturer’s instructions, with exceptions that incubation times for each solution were reduced to 3-5 min and organoids were gently agitated 10-20 times between solutions. Following dissociation (pipetted using a 200 µL tip), the 1.5 mL Eppendorf tube containing the dissociated cells was centrifuged for 5 min at 400 × g, the supernatant was removed, cells were resuspended for 15 min in 100 µL PBS on ice, and then centrifuged for 5 min at 400 × g. The supernatant was then removed, cells were resuspended in 100 µL PBS + fixable viability dye eFluor450 (eBioscience 65-0863-15, 1:1000) on ice for 15 min, centrifuged for 5 min at 400 × g, the dye solution was removed, and the cells were resuspended in 100 µL Fixing Reagent A (Life Technologies GAS002) for 15 min at room temperature (RT). Cells were then centrifuged for 5 min at 400 × g, the supernatant was removed, 10 µL Permeabilizing and Staining Reagent B with antibodies was added. Cells were incubated at RT for 15 min, then centrifuged for 5 min at 400 × g. The supernatant was then removed, and the cells washed 2× with FACS buffer. Finally, cells were resuspended in 100 µL FACS buffer and run for flow cytometry using a MAQSQuant Analyzer 10 (Miltenyi Biotech). Data were analyzed using FCS Express (DeNovo Software).

### Reagents for flow cytometry

Antibodies for flow cytometry: NeuN:PE (Novus Biologicals NBP1-92693PE); SOX2:APC (R&D Systems IC2018R-100UG); GFAP:PerCP (Novus Biologicals NBP2-33184PCP); β-tubulin III:PE:Cy7 (Biolegend 801218).

PerCP isotype control (R&D Systems IC002C). PE isotype control (Invitrogen MG2B04). PE:Cy7 isotype control (Biolegend 400232).

Compensation beads (Invitrogen 01-3333-42).

### NanoString gene expression profiling

mRNA expression was analyzed using the nCounter Human Neuropathology Panel (Catalog number: XT-CSO-HNROP1-12, NanoString), by digitally detecting and counting in a single reaction without amplification. The Neuropathology Panel consists of genes targeting six fundamental themes: neurotransmission, neuron-glia interactions, neuroplasticity, cell structure integrity, neuroinflammation, and metabolism (**Supplemental Table S1**). Each assay also includes six positive and eight negative mRNA assay controls, plus ten housekeeping mRNA controls. RNA was hybridized with the gene panel Codeset at 65 °C for 16 hr. The post-hybridization probe-target mixture was quantified using the nCounter Digital Analyzer, with subsequent data analysis performed using nSolver. Raw data were first normalized to internal positive and negative controls to eliminate variability unrelated to the samples, then normalized to the selected housekeeping genes using Geometric Means methods. Housekeeping gene expression variability of less than 20% within a single organoid was the criterion for acceptance.

mRNA extraction and preparation for NanoString was done at the COH Pathology Core. Briefly, mRNA was extracted using the RNeasy mini kit (Qiagen HB-1277-066). Concentration was measured by Nanodrop spectrophotometer ND-1000 and Qubit 3.0 Fluorometer (Thermo Scientific), with mRNA fragmentation and quality assessed by 2100 Bioanalyzer (Agilent).

NanoString data were compared against a reference database for expression of these same genes during normal cortical development (other brain regions excluded) drawn from the BrainSpan Atlas of the Developing Human Brain (https://www.brainspan.org/). To facilitate comparisons, both organoid and fetal brain datasets were normalized to the expression of the same NanoString internal reference genes (**Supplemental Table S1**) using *Singscore* ^32^. *Singscore* ranks gene expression and calculates a non-parametric score for individual genes in each gene set (NanoString or BrainSpan). This rank-based approach is robust to differences in the underlying data distribution, enabling scoring of gene sets across different transcriptomic datasets and assessing gene set activity in individual samples. In the single-sample presentation employed here, scores range between 0.0 and 1.0 and represent the extent to which each gene in the gene set is expressed as compared to the distribution of reference genes, and scored as higher (score > 0.5) or lower (score < 0.5) (**Supplemental Figure S2**).

### Statistical comparisons

Tests for statistical significance (*p* < 0.05), and internal correlations of NanoString and BrainSpan data (**Figure 6**), were made using Prism v9 (Graph Pad).

### Gene Ontology

Tissue integrity genes presented in **Figure 1F** were interrogated for gene ontology (https://geneontology.org) ^64,65^ and compared using the “Gene ontology enrichment analysis for biological processes” tool ^66^.

## ACKNOWLEDGEMENTS

The authors acknowledge and appreciate the scientific discussions and technical assistance of Drs. Brian Armstrong and Yate-Ching Yuan, and the editorial assistance of Andrea Lynch, MLIS, and Drs. Chris Gandhi, Erin Keebaugh and Keely Walker in the Office of Faculty and Institutional Support.

This work was supported in part by National Institutes of Health grant P30CA033572 and its Developmental Cancer Therapeutics program, and by the Albert & Bettie Sacchi Foundation. Research reported in this publication includes work performed in the Research Pathology Services Core, the Light Microscopy & Digital Imaging Shared Resource, the High Throughput Screening Core, and Bioinformatics Core in the Division of Translational Bioinformatics, all supported by the National Cancer Institute of the NIH under grant number P30CA033572. The content is solely the responsibility of the authors and does not necessarily represent the official views of the National Institutes of Health.

## AUTHOR CONTRIBUTIONS

Conceptualization: A.V.O., M.G., M.E.B.

Methodology: A.V.O., B.B., N.C., M.G., M.E.B.

Software Programming: n/a

Validation: n/a

Formal Analysis: R.R., D.O’M., H.H.Y., M.E.B.

Investigation: A.V.O., D.A.

Resources: C.E.B., M.G.

Data Curation: D.A., B.B., M.E.B.

Writing – Original Draft: A.V.O., D.A. M.E.B.

Writing – Review and Editing: A.V.O., D.A., M.G., M.E.B.

Visualization: A.V.O., D.A., M.E.B.

Supervision: M.G., M.E.B.

Project Administration: n/a

Funding Acquisition: C.E.B., M.G., M.E.B.

## DECLARATION OF INTERESTS

M.G. is a founder and holds equity in StemCellMed Inc., a firm utilizing L-*MYC*-immortalized NSCs and their derivatives. The other authors declare no relevant competing interests.

