## Supplemental Figure 1 for "Modeling cerebral development *in vitro* with L-*MYC*-immortalized human neural stem cell-derived organoids"

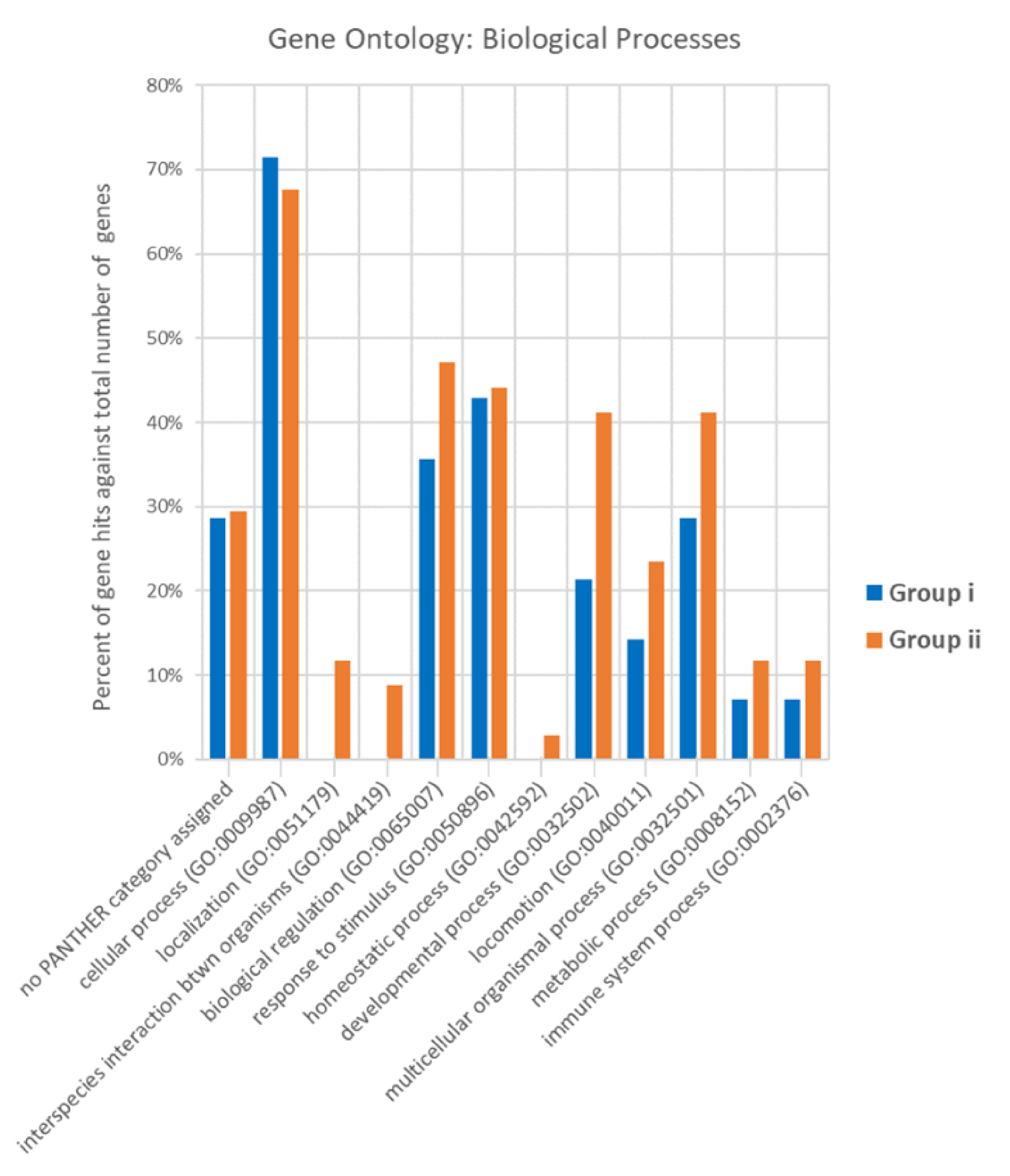


**Supplemental Figure S1**

Gene ontology analysis of genes in the Tissue Integrity theme of NanoString, separated into two groups as described in the text.
