## Supplemental Figure 2 for "Modeling cerebral development *in vitro* with L-*MYC*-immortalized human neural stem cell-derived organoids"

### SUPPLEMENTAL FIGURE S2

#### ORGANOIDS (NANOSTRING) AND CORTEX (BRAINSPAN): GRAPHS OF CELL TYPES, PATHWAYS AND THEMES

Plots of expression data (as SingScores) for developing organoids and fetal cortex often overlap in magnitude and waveform.

Graphs of expression data from organoids (NanoString; **above**) and fetal cortex (BrainSpan; **below**), presented as SingScores. SingScores vary between 0.0 and 1.0, and represent the extent to which each gene in the gene set (organoid or cortex) is expressed as compared to a common set of reference genes. Expression relative to the set of reference genes is scored as higher (score > 0.5) or lower (score < 0.5). The genes defining each tissue or process, and the reference genes, are listed in **Supplemental Table 1**.

### Group 1: Cell Types

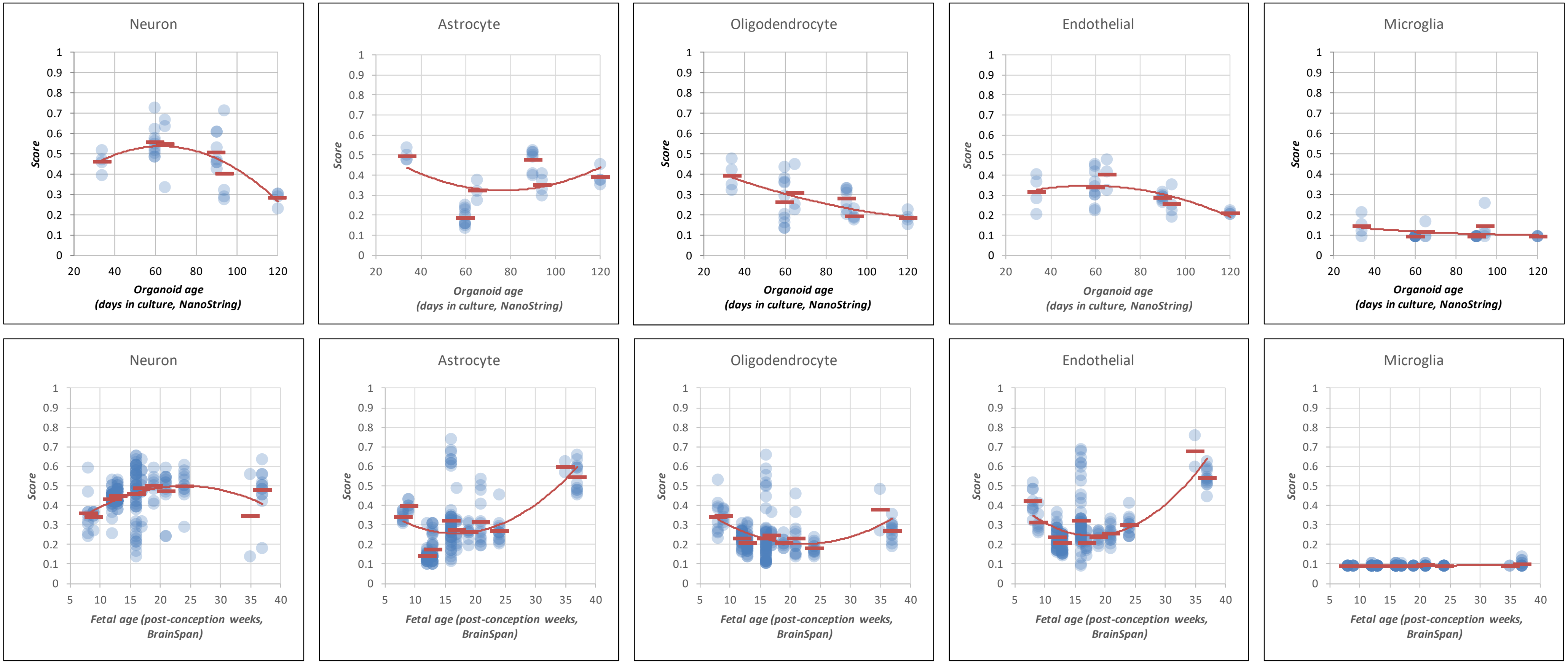

### Group 2: Pathways and Themes

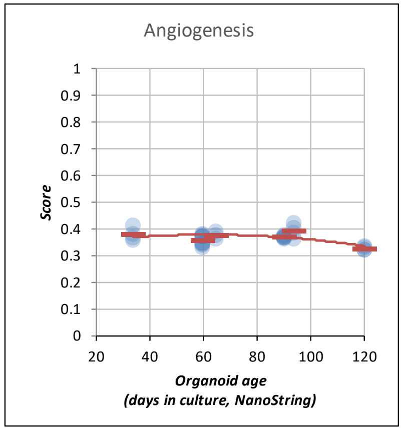


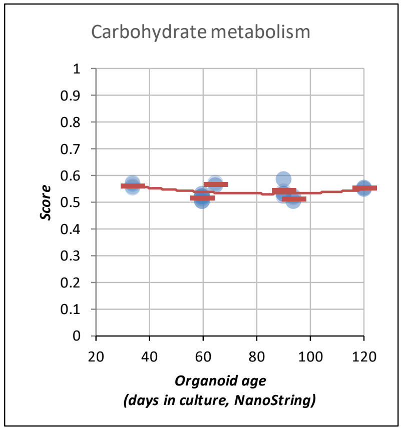


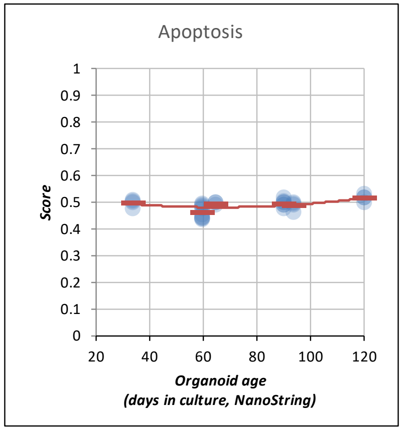


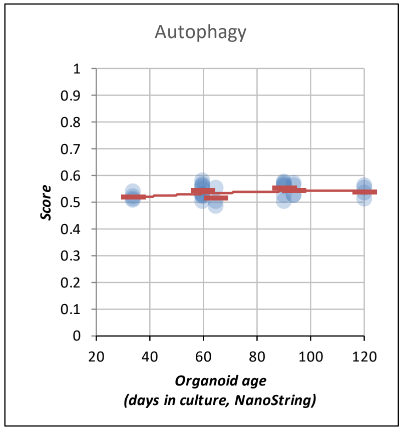


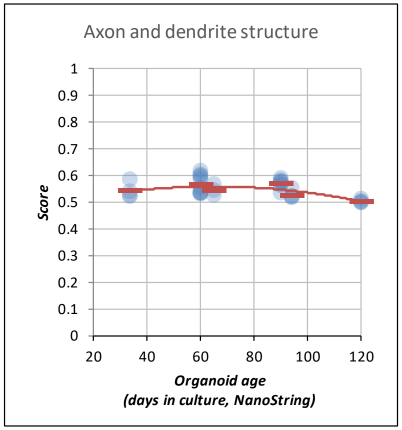


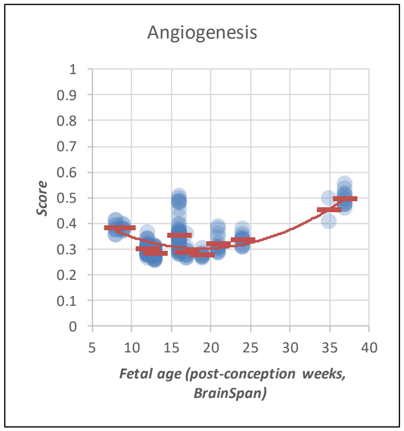


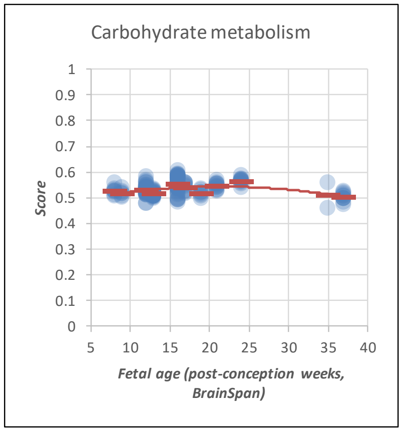


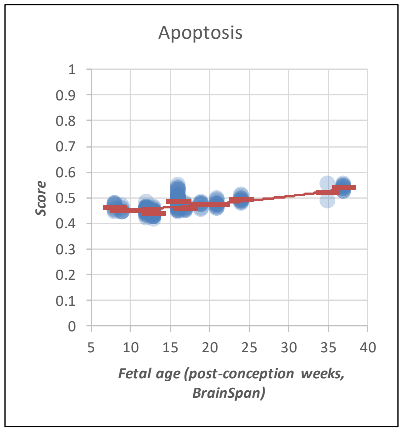


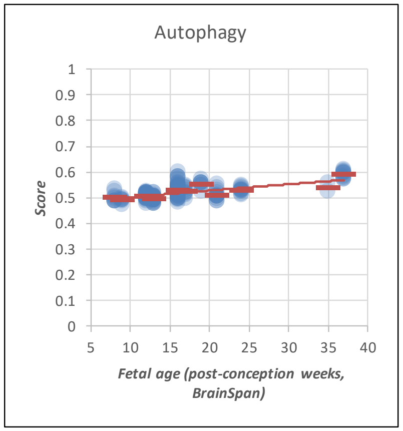


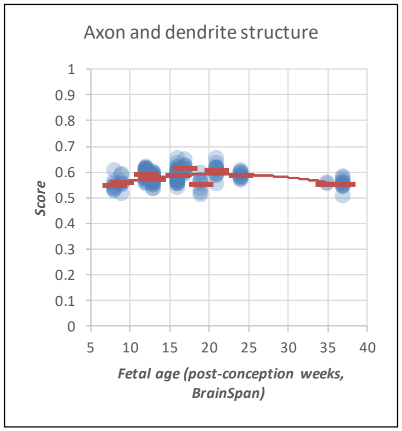

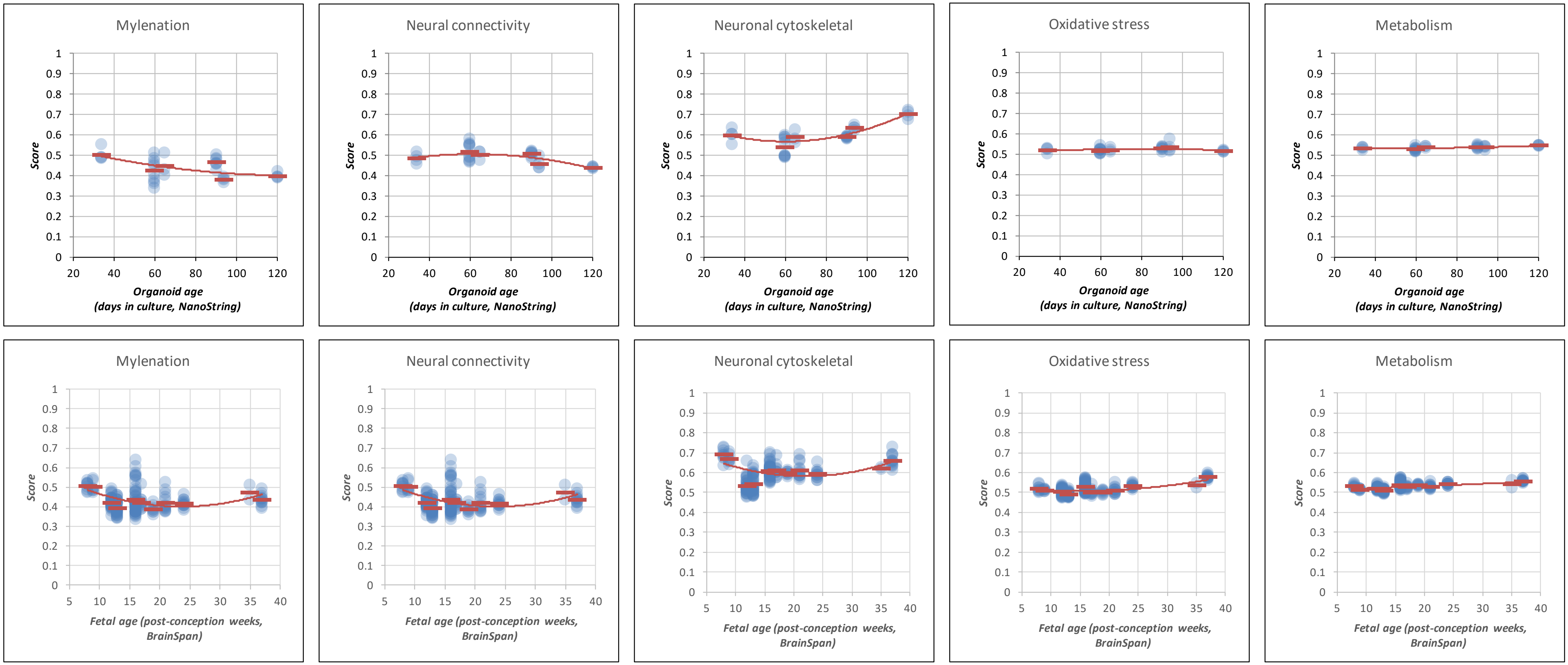
Mylenation
Neural connectivity
Neuronal cytoskeletal
Oxidative stress
Metabolism
Score
Organoid age
(days in culture, NanoString)
Mylenation
Neural connectivity
Neuronal cytoskeletal
Oxidative stress
Metabolism
Score
Fetal age (post-conception weeks, BrainSpan)

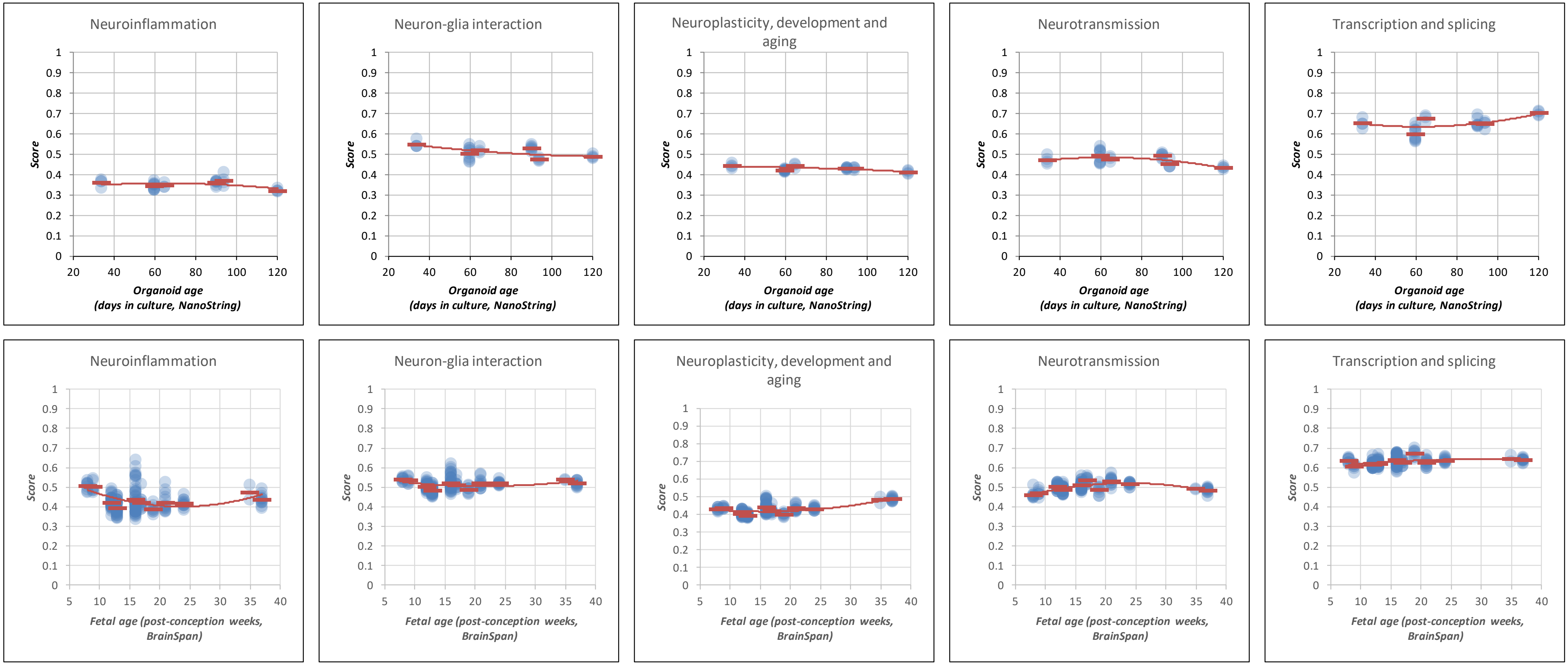
Neuroinflammation
Neuron-glia interaction
Neuroplasticity, development and aging
Neurotransmission
Transcription and splicing
Score
Organoid age
(days in culture, NanoString)
Neuroinflammation
Neuron-glia interaction
Neuroplasticity, development and aging
Neurotransmission
Transcription and splicing
Score
Fetal age (post-conception weeks, BrainSpan)

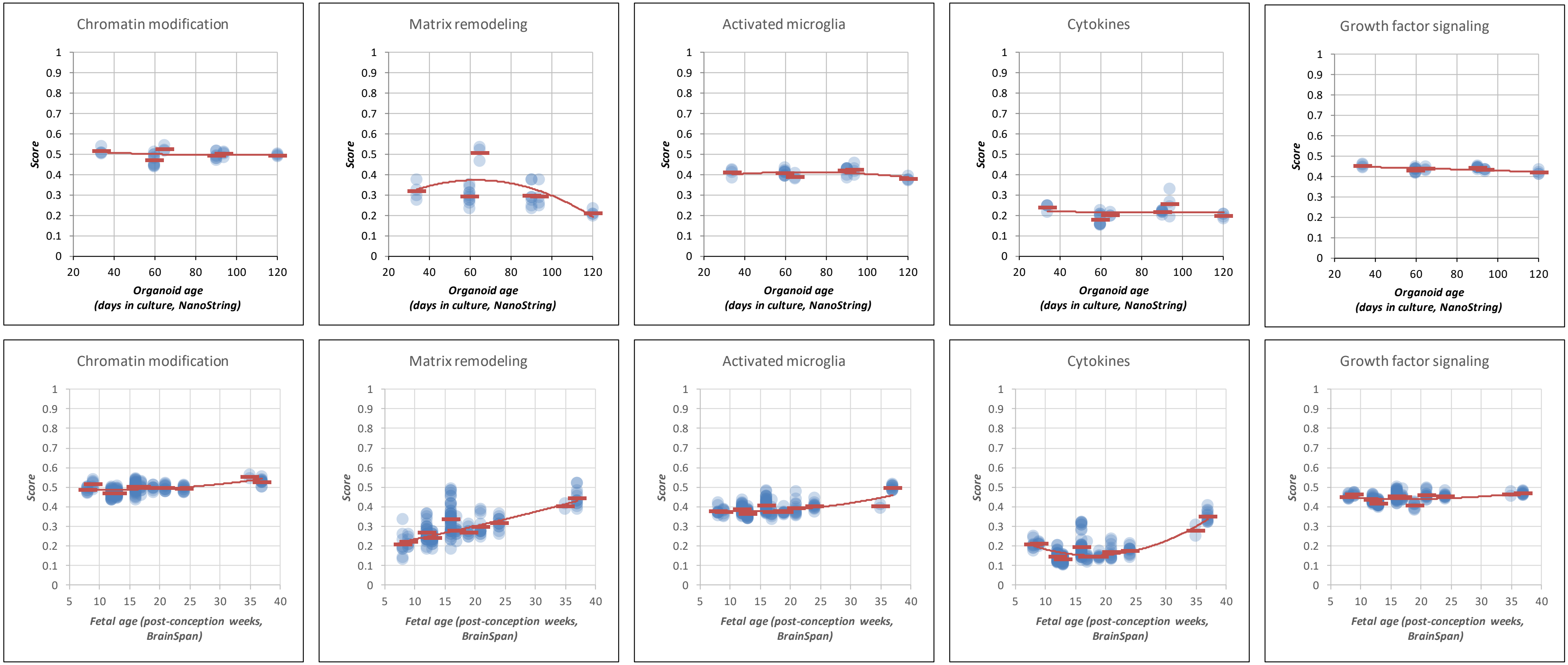
Chromatin modification
Matrix remodeling
Activated microglia
Cytokines
Growth factor signaling
Score
Organoid age
(days in culture, NanoString)
20 40 60 80 100 120
0 0.1 0.2 0.3 0.4 0.5 0.6 0.7 0.8 0.9 1
Chromatin modification
Matrix remodeling
Activated microglia
Cytokines
Growth factor signaling
Score
Fetal age (post-conception weeks, BrainSpan)
5 10 15 20 25 30 35 40
0 0.1 0.2 0.3 0.4 0.5 0.6 0.7 0.8 0.9 1

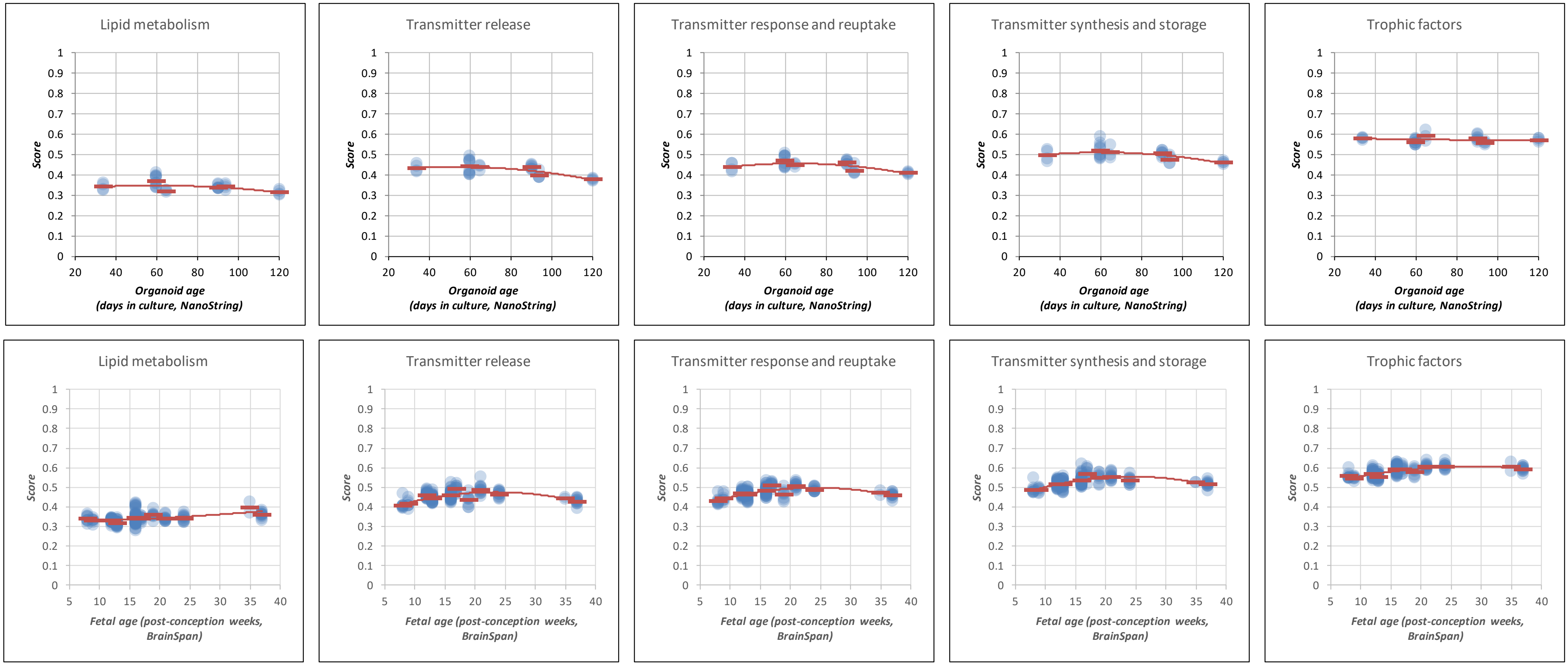
Lipid metabolism
Transmitter release
Transmitter response and reuptake
Transmitter synthesis and storage
Trophic factors
Score
Organoid age
(days in culture, NanoString)
Lipid metabolism
Transmitter release
Transmitter response and reuptake
Transmitter synthesis and storage
Trophic factors
Score
Fetal age (post-conception weeks, BrainSpan)

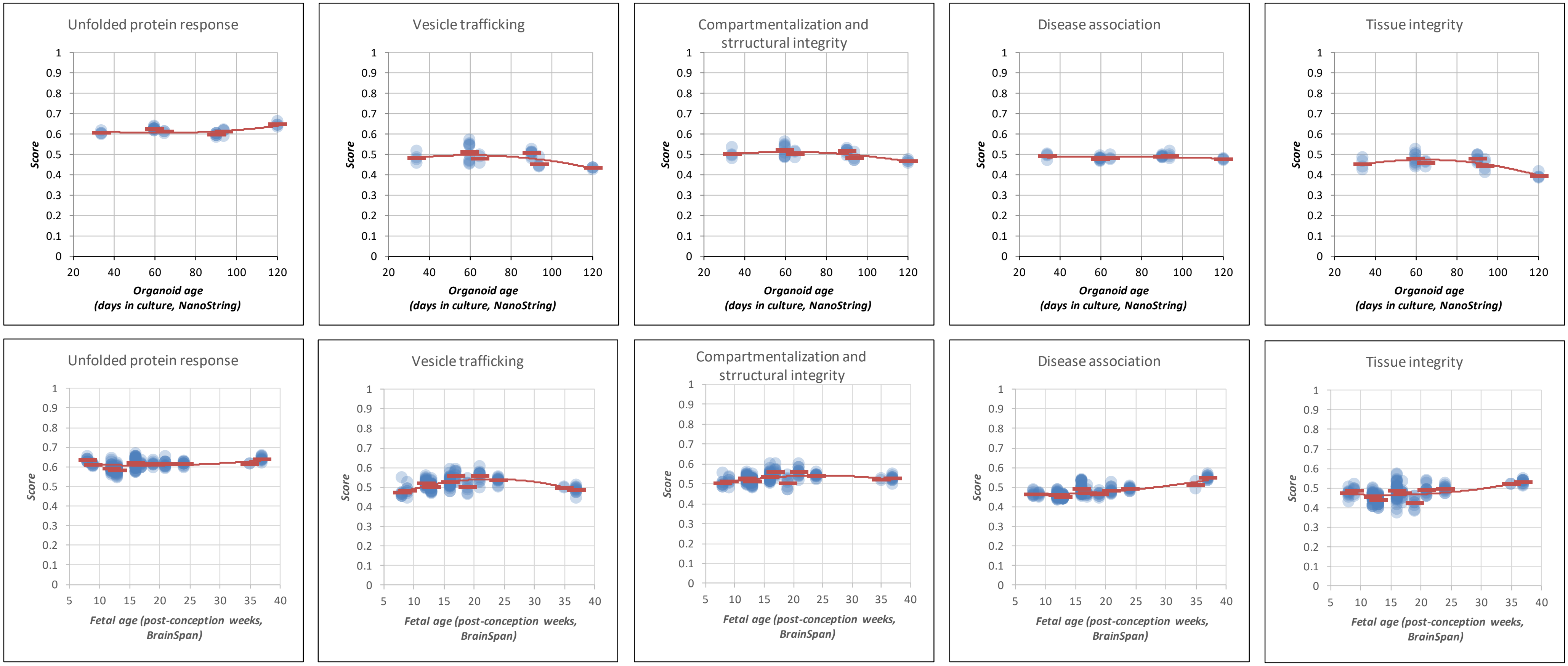
Unfolded protein response
Vesicle trafficking
Compartmentalization and strructural integrity
Disease association
Tissue integrity
Score
Organoid age (days in culture, NanoString)
Unfolded protein response
Vesicle trafficking
Compartmentalization and strructural integrity
Disease association
Tissue integrity
Score
Fetal age (post-conception weeks, BrainSpan)

### Group 3: Reference Genes

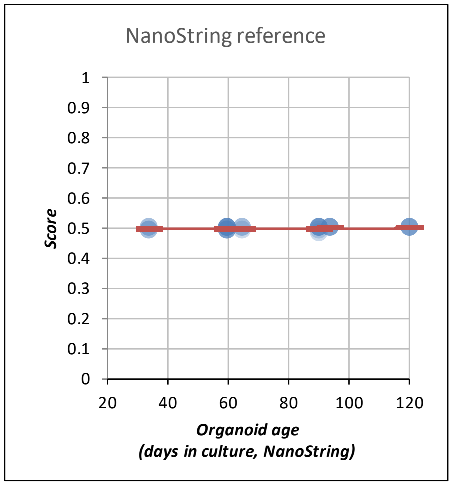


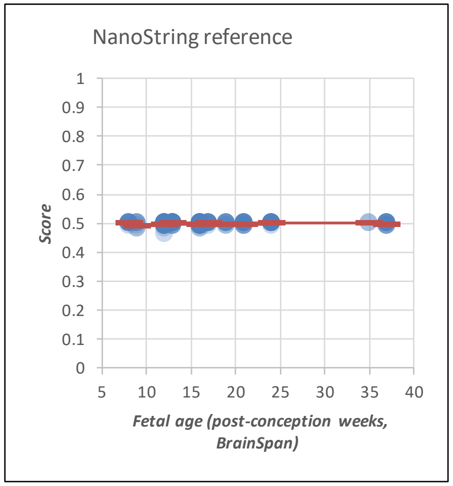
