## Supplemental Table S1 for "Modeling cerebral development *in vitro* with L-*MYC*-immortalized human neural stem cell-derived organoids"

### GENE LISTS DEFINING CELL TYPES, PATHWAYS AND THEMES (NANOSTRING)

#### Cell Types

| Astrocytes (8) | Endothelial Cells (9) | Microglia (12) | Neurons (6) | Oligodendrocytes (13) |
|---|---|---|---|---|
| ALDH1L1 | CLDN5 | AIF1 | DLX1 | BCAS1 |
| EGFR | EMCN | CX3CR1 | DLX2 | ERBB3 |
| ENTPD2 | ESAM | GPR84 | GRM2 | FA2H |
| GDPD2 | FLT1 | IRF8 | ISLR2 | GAL3ST1 |
| ITGA7 | ICAM2 | ITGAM | SLC17A6 | GJB1 |
| KIAA1161 | LSR | LRRC25 | TBR1 | GSN |
| NWD1 | MYCT1 | NCF1 | | MYRF |
| SOX9 | NOSTRIN | P2RY12 | | NINJ2 |
| | TIE1 | SPI1 | | PLLP |
| | | TLR2 | | PLXNB3 |
| | | TMEM119 | | PRKCQ |
| | | TNF | | SOX10 |
| | | | | UGT8 |

#### Pathways

| Activated Microglia (92) | Angiogenesis (78) | Apoptosis (61) | Autophagy (33) | Axon and Dendrite Structure (160) | Carbohydrate Metabolism (44) | Chromatin Modification (62) | Cytokines (52) | Disease Association (307) | Growth Factor Signaling (150) | Lipid Metabolism (41) | Matrix Remodeling (7) | Myelination (47) | Neural Connectivity (166) | Neuronal Cytoskeleton (17) | Oxidative Stress (91) | Tissue Integrity (45) | Transcription and Splicing (47) | Transmitter Release (165) | Transmitter Response and Reuptake (148) | Transmitter Synthesis and Storage (59) | Trophic Factors (48) | Unfolded Protein Response (48) | Vesicle Trafficking (156) |
|---|---|---|---|---|---|---|---|---|---|---|---|---|---|---|---|---|---|---|---|---|---|---|---|
| ADCY8 | ACVRL1 | AKT1 | AP1S1 | ACVRL1 | AKT1 | ATG5 | CCL2 | ABAT | | ACAA1 | MMP12 | AKT1 | ACHE | ACTN1 | ABL1 | AQP4 | ACIN1 | ABAT | ADCY5 | ABAT | ABL1 | ATF4 | ABAT |
| ADRA2A | ADRB2 | AKT2 | AP3M2 | ADCY9 | AKT1S1 | ATM | CCL5 | ACHE | AKT2 | ACHE | MMP14 | AKT2 | ADAM10 | DES | AGER | CADM3 | ATXN2 | ACHE | ADCY8 | ACHE | AKT1 | ATF6 | ACHE |
| AMIGO1 | ANG | AKT3 | AP3S1 | ADCYAP1 | AKT2 | ATXN3 | CCR2 | ACVRL1 | AKT3 | ARSA | MMP16 | AMIGO1 | ADCYAP1 | DNAH1 | AIF1 | CD34 | BCAS2 | ADCY5 | ADCY9 | ADCYAP1 | AKT2 | ATP6V0D1 | ADCYAP1 |
| AP3M2 | ANGPT2 | APAF1 | AP4S1 | ADORA1 | AKT3 | ATXN7 | CCR5 | ADAM10 | APC | B4GALT6 | MMP19 | ARHGEF10 | ADORA1 | GFAP | AKT1 | CD4 | CCNH | ADCY8 | ADORA1 | AP2A2 | AKT3 | ATXN3 | ADORA1 |
| APOE | C3 | ATF4 | ARSA | ADRB2 | BCHE | BRMS1L | CD40 | ADCY5 | ARRB2 | CDS1 | MMP2 | ATRN | ADRA2A | GSN | APOE | CD40 | CDC40 | ADCY9 | ADORA2A | AP2B1 | ATF4 | BAX | ADRB2 |
| ARRB2 | C5 | ATM | ATP6V0C | AGER | CAB39 | CAMK2D | CNTF | ADCYAP1 | ATF4 | CERS1 | MMP24 | CD9 | ADRB2 | INA | APP | CD8A | CDK7 | ADORA2A | ADRA2A | ATP6V0C | BAD | BCL2 | AKT1 |
| ATP6V0E1 | C6 | BAD | ATP6V0D1 | AMIGO1 | CCND1 | CDK2 | CSF1 | ADORA2A | ATM | CERS2 | MMP9 | CLDN5 | AKT1 | KATNA1 | ATF4 | CLDN15 | DDX23 | ADRB2 | ADRB2 | ATP6V0D1 | BAX | CAST | AMPH |
| ATP6V1A | CCL2 | BAX | ATP6V1H | AP1S1 | CPT1B | CHD4 | CSF1R | ADRA2A | BCL2 | CERS4 | | CLU | AMPH | MAPT | ATP13A2 | CLDN5 | GTF2A1 | AKT1 | AKT1 | ATP6V0E1 | BCL2 | CCL2 | APOE |
| ATP8A2 | CCL5 | BCL2 | C9orf72 | AP3M2 | CREB1 | CREBBP | CSF2RB | AGER | BDNF | CERS6 | | CNTNAP1 | AP1S1 | MSN | ATP7A | CNTN1 | GTF2B | AKT2 | AKT2 | ATP6V0E2 | BDNF | CCND1 | ARC |
| BCL2 | CCR2 | BCL2L1 | CD68 | AP3S1 | CRTC2 | CRTC2 | CX3CL1 | AIF1 | CACNA1A | CHAT | | CTNNB1 | APC | NEFH | ATRN | CNTNAP1 | GTF2H1 | AKT3 | AKT3 | ATP6V1A | CALM1 | CDC27 | ARRB2 |
| C3 | CD34 | BID | CHMP2B | APC | DLAT | CTNNB1 | CX3CR1 | AKT1 | CACNA1B | DGKB | | CXCR4 | APP | NEFL | BAD | CNTNAP2 | GTF2H3 | APP | ATF4 | ATP6V1D | CALML5 | CDK5RAP3 | ATXN3 |
| CASP8 | CHRNA7 | CASP1 | CLN3 | APOE | DLD | CXXC1 | CXCL10 | ANG | CACNA1C | DGKE | | EGR2 | ARC | NES | BCL2 | EFNA1 | GTF2IRD1 | ARRB2 | ATP2B3 | ATP6V1E1 | CAMK2B | CUL1 | BCHE |
| CCL5 | COL4A1 | CASP3 | CTNS | APP | GGT1 | DOT1L | CXCL11 | ANGPT2 | CACNA1D | GAL3ST1 | | FA2H | ARHGAP44 | PFN1 | BNIP3 | EFNA5 | HNRNPM | ATF4 | BAD | ATP6V1G2 | CAMK2D | CUL2 | CA2 |
| CCR2 | COL4A2 | CASP6 | CTSE | ARC | GLS | EHMT1 | CXCL12 | AP2A2 | CACNA1F | GALC | | FAM126A | ARRB2 | PLS1 | CASP3 | EFNB3 | HSPA6 | ATP2B3 | BDNF | ATP6V1H | CAMK2G | CUL3 | CACNA1A |
| CCR5 | CSPG4 | CASP7 | ENTPD4 | ARHGAP44 | GSK3B | EP300 | CXCL16 | AP2B1 | CACNA1S | GBA | | GAL3ST1 | ATCAY | RDX | CCL5 | EPHA2 | LSM2 | BCL2 | CACNA1A | ATP7A | CAMK4 | CXXC1 | CACNA1B |
| CD33 | CTNNB1 | CASP8 | GAA | ARRB2 | GSS | ERG | CXCR4 | AP3M2 | CACNB2 | GDPD2 | | GJB1 | ATP6V0D1 | SPTBN2 | CCS | EPHA3 | LSM7 | CACNA1A | CACNA1B | CACNA1A | DLX1 | DDIT3 | CACNB2 |
| CD44 | CX3CL1 | CASP9 | GALC | ATCAY | IDE | FMR1 | EGF | APAF1 | CACNB4 | GGT1 | | HEXB | BCAS1 | TPM1 | CDK2 | EPHA4 | MNAT1 | CACNA1B | CACNA1C | CACNA1B | DLX2 | DNAJA2 | CACNB4 |
| CD68 | CX3CR1 | CDKN1A | GBA | AVP | IDH1 | GATA2 | EGFR | APOE | CALM1 | GPD1L | | HGF | BCL2L1 | | CLN8 | EPHA5 | PCNA | CACNA1C | CACNA1D | CHAT | FASLG | EIF2S1 | CADPS |
| CD9 | CXCL10 | CSF2RB | GGA1 | BACE1 | IGF1 | HDAC1 | EPO | APP | CAMK2B | HPGDS | | JAM3 | CACNA1A | | CYBB | EPHA6 | POLR2B | CACNA1D | CACNA1F | CHRNB2 | GSK3B | EP300 | CALB1 |
| CDS1 | DLL4 | CYCS | GNPTAB | BECN1 | IGF1R | HDAC2 | FAS | ARHGEF10 | CAMK2D | LCLAT1 | | KCNJ10 | CACNA1B | | CYCS | EPHA7 | POLR2H | CACNA1F | CACNA1S | COMT | HRAS | ERLEC1 | CALM1 |
| CLU | EFNA1 | DDIT3 | GNPTG | BNIP3 | INS | HDAC6 | FASLG | ATF4 | CAMK2G | LSR | | KEL | CACNA1C | | DDIT3 | ESAM | POLR2J | CACNA1S | CALM1 | CPLX1 | IKBKB | GFPT1 | CAMK2B |
| CR1 | EGF | EIF2S1 | GUSB | CA2 | INSR | HDAC7 | FLT1 | ATF6 | CASP3 | LYPLA1 | | KLK6 | CACNB4 | | DNM2 | FYN | POLR2K | CACNB2 | CALML5 | DAGLA | JUN | HSPA6 | CCL2 |
| CSF1 | EGFL7 | FAS | HEXB | CACNA1A | LDHC | IL1B | FLT4 | ATM | CCND1 | PLA2G16 | | LAMA2 | CADPS | | EP300 | HLA-DRA | POLR2L | CACNB4 | CAMK2B | DBH | KRAS | HSPB1 | CDK5 |
| CSF1R | EMCN | FASLG | LAMP1 | CACNA1B | MTOR | LIF | HGF | ATP6V0C | CD14 | PLA2G2A | | LPAR1 | CALB1 | | FAS | ICAM1 | PQBP1 | CALM1 | CAMK2D | DDC | MAP2K1 | KEAP1 | CHAT |
| CSPG4 | EMP2 | FOS | MAN2B1 | CALB1 | NPY | MAPK3 | IL10 | ATP6V0D1 | CDS1 | PLA2G2E | | MAG | CALB2 | | FN1 | ICAM2 | PRPF3 | CALML5 | CAMK2G | DNM2 | MAP2K2 | LMNA | CHRM5 |
| CTSE | ENG | HRAS | NAGLU | CALB2 | PDPK1 | MAPK8 | IL10RA | ATP6V1D | CHRM5 | PLA2G2F | | MAL | CAMK2B | | FOS | ISLR2 | PRPF31 | CAMK2B | CAMK4 | DRD1 | MAPK1 | MAPK10 | CHRNA7 |
| CUL1 | EPHA2 | HTRA2 | NAPSA | CAMK2B | PGAM1 | MEAF6 | IL13RA1 | ATP6V1E1 | CNTF | PLA2G4A | | MBP | CAMK2D | | FXN | ITGA7 | RAN | CAMK2D | CHRM5 | DRD2 | MAPK10 | MAPK8 | CHRNB2 |
| CX3CR1 | FASLG | IKBKB | NPC1 | CAMK2D | PGK1 | MECP2 | IL15RA | ATXN2 | CREB1 | PLA2G4B | | MOG | CCK | | GNAO1 | ITGAL | SART1 | CAMK2G | CHRNA7 | DRD4 | MAPK3 | MAPK9 | CNR1 |
| CXCL10 | FGF2 | ITPR1 | NPC2 | CCK | PIK3CA | MTA1 | IL1B | BACE1 | CREBBP | PLA2G4C | | MTOR | CCL2 | | GPR37 | ITGAM | SF3A2 | CAMK4 | CHRNB2 | GAD1 | MAPK8 | MFN2 | CNTN4 |
| CXCL16 | FLT1 | ITPR2 | PPT1 | CCL2 | PIK3CB | MTA2 | IL1R1 | BAD | CSF1R | PLA2G4D | | MYRF | CDK5 | | GSK3B | JAM3 | SF3B2 | CASP3 | CNR1 | GAD2 | MAPK9 | NFE2L2 | CPLX1 |
| CXCR4 | FLT4 | ITPR3 | SLC11A1 | CCR2 | PIK3R1 | MTHFR | IL4R | BAX | CTNNB1 | PLA2G4E | | NF1 | CDK5R1 | | GSR | L1CAM | SF3B4 | CDK5 | CREB1 | GPR37 | MAPKAPK2 | PIK3R1 | CREB1 |
| CYBB | FN1 | JUN | SQSTM1 | CDK5 | PPARGC1A | NELFA | IL6 | BCHE | CUL1 | PLA2G4F | | NGFR | CHAT | | GSS | LRRC4 | SLU7 | CHAT | CREBBP | GRIN2A | NFKBIA | PLA2G4B | CRH |
| DNM1L | GATA2 | KRAS | TCIRG1 | CDK5R1 | PPP2CA | NOS1 | IL6R | BCL2 | CUL3 | PLA2G6 | | NKX6-2 | CHRM5 | | GSTP1 | MAG | SMN1 | CHRM5 | CRH | HTR1A | NFKBIB | PRKCSH | CTNNB1 |
| EGR2 | GPR4 | LMNA | | CHL1 | PPP2R5C | PCNA | INHBB | BCL2L1 | DDIT3 | PTDSS1 | | NRG1 | CHRNA7 | | GUCY1B3 | MPZ | SNRPA | CHRNA7 | DAGLA | NOS1 | NGF | PRKN | CX3CL1 |
| ENTPD4 | HDAC7 | MAP2K1 | | CLDN5 | PPP2R5E | PHF19 | LIF | BDNF | DLL4 | PTDSS2 | | NTF3 | CHRNB2 | | HDAC2 | NCAM1 | SRSF4 | CHRNB2 | DLG4 | NR4A2 | NGFR | PSMB8 | CX3CR1 |
| FLT1 | HGF | MAP2K2 | | CLU | PRKAA2 | PHF2 | LTBR | BID | DRD2 | PTGS2 | | OLIG2 | CLN3 | | HDAC6 | NEGR1 | TAF10 | COMT | DRD1 | NSF | NTF3 | PSMB9 | DBH |
| FN1 | HIF1A | MAPK1 | | CNR1 | RAB2A | PHF21A | NGFR | C3 | EFNA1 | SGPL1 | | PLLP | CLN8 | | HGF | NEO1 | TAF4 | CREB1 | DRD2 | PAH | NTRK1 | RAD23B | DLG3 |
| GAA | HMGB1 | MAPK10 | | CNTF | SIRT1 | PKN1 | OSMR | C5 | EFNA5 | SMPD4 | | PMP22 | CNKSR2 | | HIF1A | NINJ2 | TAF4B | CRH | DRD4 | PARK7 | PDPK1 | SEC23A | DLG4 |
| GALC | HMOX1 | MAPK3 | | CNTN1 | SLC1A1 | PPARGC1A | PDGFRB | C6 | EFNB3 | TAZ | | PTEN | COMT | | HMOX1 | NLGN4X | TAF6L | CYP4X1 | ENTPD2 | PDE1B | PIK3CA | TRIM37 | DLGAP1 |
| GBA | HSPB1 | MAPK8 | | CNTN4 | SLC1A2 | PRKAA2 | PLEKHO2 | C9orf72 | EFR3A | UGCG | | SCN2A | CPLX1 | | HSPB1 | NRXN1 | TAF9 | DDC | EP300 | PNKD | PIK3CB | UBE2K | DRD1 |
| GNG2 | IL1B | MAPK9 | | CNTNAP1 | SOD1 | PRKCA | PRL | CA2 | EGF | UGT8 | | SH3TC2 | CTNNB1 | | HTRA2 | NTNG1 | TARDBP | DLG4 | F2 | RAB3A | PIK3R1 | UBE2N | DRD2 |
| GNPTAB | IL6 | NFKBIA | | CNTNAP2 | SUCLA2 | PRKCB | TGFB1 | CACNA1C | EGFR | | | SIRT2 | CX3CL1 | | IDH1 | PECAM1 | TBP | DLGAP1 | FOS | RIMS1 | PSEN1 | UBE3A | DRD4 |
| GNPTG | ITGA5 | NGF | | COMT | TFAM | RING1 | TGFBR2 | CACNA1D | EGR1 | | | SOD1 | DDC | | IL1B | SLC2A1 | TBPL1 | DRD1 | GABRA1 | SLC17A6 | PSEN2 | UBQLN1 | EGR2 |
| GPR84 | JAM3 | NTRK1 | | CPLX1 | VIP | SIRT1 | TNF | CACNA1F | EP300 | | | TGFB1 | DES | | IL1R1 | SRC | TCERG1 | DRD2 | GABRA4 | SLC18A2 | RAC1 | USP30 | FGF12 |
| GPRASP1 | JUN | PARP1 | | CREB1 | | SIRT2 | TNFRSF10A | CACNA1S | EPHA2 | | | UGT8 | DLG3 | | IL6 | TNC | TWISTNB | DRD4 | GABRB2 | SLC18A3 | RAF1 | VCP | FGF14 |
| GRN | LIF | PDPK1 | | CRH | | SIRT7 | TNFRSF10B | CADPS | EPHA3 | | | XK | DLG4 | | INS | | U2AF2 | EGFR | GABRB3 | SLC32A1 | RELA | WFS1 | FMR1 |
| GUSB | MMP14 | PIK3CA | | CTNNB1 | | SMYD1 | TNFRSF10D | CALB1 | EPHA4 | | | ZNF24 | DLGAP1 | | IPCEF1 | | XAB2 | ERBB3 | GABRG2 | SLC6A3 | RHOA | XBP1 | GABRA1 |
| HIF1A | MMP19 | PIK3CB | | DAGLA | | SNCA | TNFRSF11B | CALM1 | EPHA5 | | | | DNM1L | | JUN | | | FOS | GABRP | SLC6A4 | TP53 | XIAP | GABRA4 |
| IGF1 | MMP2 | PIK3R1 | | DCX | | SOX10 | TNFRSF12A | CALML5 | EPHA6 | | | | DNM2 | | LPO | | | FYN | GABRR1 | SNCA | | | GABRB2 |
| IL1B | MMRN2 | PRF1 | | DDC | | SP1 | TNFRSF1A | CAMK2G | ERBB3 | | | | DRD1 | | LRRK2 | | | GABRA1 | GABRR3 | SNCAIP | | | GABRB3 |
| IL1R1 | NCL | RAF1 | | DLG3 | | SPI1 | TNFRSF1B | CASP1 | FAM126A | | | | DRD2 | | MAPT | | | GABRA4 | GLRB | SNCB | | | GABRG2 |
| IL6R | NF1 | RELA | | DLG4 | | TAF10 | VEGFA | CASP3 | FAS | | | | DRD4 | | MGMT | | | GABRB2 | GNAI1 | STX1A | | | GABRR1 |
| IRF8 | NOS3 | TLR4 | | DRD1 | | TAF6L | | CASP7 | FASLG | | | | EPHA4 | | MMP14 | | | GABRB3 | GNAI2 | STX1B | | | GABRR3 |
| ITGA5 | NOTCH1 | TNF | | DRD2 | | TAF9 | | CASP8 | FGF12 | | | | EPHA7 | | MUTYH | | | GABRG2 | GNAI3 | STX2 | | | GAD1 |
| ITGAL | NOTCH3 | TNFRSF10A | | DRD4 | | TGFB1 | | CASP9 | FGF14 | | | | FMR1 | | NCF1 | | | GABRP | GNAO1 | SYT1 | | | GAD2 |
| ITGAX | NOTCH4 | TNFRSF10B | | EPHA4 | | TP53 | | CAST | FGF2 | | | | FRMPD4 | | NEFH | | | GABRR1 | GNAQ | TCIRG1 | | | GDNF |
| JAM3 | NRXN1 | TNFRSF10D | | EPHA5 | | TRIM28 | | CCL2 | FOS | | | | FUS | | NFE2L2 | | | GABRR3 | GNB5 | TH | | | GFAP |
| LMNA | NTRK1 | TNFRSF1A | | EPHA7 | | TRIM37 | | CCL5 | FYN | | | | FYN | | NGFR | | | GAD1 | GNG2 | UNC13A | | | GLRB |
| LOX | PDGFRB | TP53 | | FAS | | UBE2N | | CCND1 | GDNF | | | | GABRA1 | | NME5 | | | GAD2 | GNGT1 | VIP | | | GLS |
| MARCO | PIK3CA | TRADD | | FMR1 | | USP21 | | CCR2 | GSK3B | | | | GABRA4 | | NOL3 | | | GDNF | GRIA1 | | | | GNAI1 |
| MMP12 | PIK3CB | XIAP | | FRMPD4 | | VEGFA | | CCR5 | HDAC1 | | | | GABRB2 | | NOS1 | | | GLRB | GRIA2 | | | | GNAI2 |
| MMP14 | PRKCA | | | FUS | | XBP1 | | CCS | HDAC2 | | | | GABRB3 | | NOS3 | | | GLS | GRIA3 | | | | GRIA1 |
| NAGLU | PRKCB | | | GABRG2 | | | | CD14 | HGF | | | | GABRG2 | | NQO1 | | | GNAI1 | GRIA4 | | | | GRIA2 |
| NLRP3 | PTEN | | | GAD2 | | | | CD33 | HRAS | | | | GABRP | | NR4A2 | | | GNAI2 | GRIK2 | | | | GRIA3 |
| NPAS4 | PTGS2 | | | GLRB | | | | CD4 | HSPA6 | | | | GABRR1 | | OXR1 | | | GNAI3 | GRIN1 | | | | GRIA4 |
| NPC2 | RHOA | | | GNAI2 | | | | CD44 | HSPB1 | | | | GABRR3 | | PARK7 | | | GNAO1 | GRIN2A | | | | GRIK2 |
| OPTN | RRAS | | | GNAO1 | | | | CD68 | IGF1 | | | | GAD1 | | PARP1 | | | GNAQ | GRIN2B | | | | GRIN1 |
| P2RX4 | SHH | | | GNAQ | | | | CD9 | IKBKB | | | | GAD2 | | PDGFRB | | | GNB5 | GRIN2C | | | | GRIN2A |
| P2RY12 | SIRT1 | | | GRIA1 | | | | CDK2 | IL1B | | | | GLRB | | PINK1 | | | GNG2 | GRIN2D | | | | GRIN2B |
| PGK1 | SP100 | | | GRIA3 | | | | CDK5 | IL1R1 | | | | GRIA1 | | PLA2G4A | | | GNGT1 | GRIN3B | | | | GRIN2C |
| PLCL2 | STAB1 | | | GRIA4 | | | | CDK5R1 | IL6 | | | | GRIA2 | | PPARGC1A | | | GRIA1 | GRM1 | | | | GRIN2D |
| PLXNC1 | STAT1 | | | GRIK2 | | | | CDK5RAP3 | INPP4A | | | | GRIA3 | | PRNP | | | GRIA2 | GRM2 | | | | GRIN3B |

| | | | | | | | | | | | | | | | | | | | | | | | |
|---|---|---|---|---|---|---|---|---|---|---|---|---|---|---|---|---|---|---|---|---|---|---|---|
| PPM1L | TGFBR2 | | | GRIN1 | | | | CDK7 | INPP5F | | | | GRIA4 | | PSEN1 | | | GRIA3 | GRM5 | | | | GRM1 |
| PSEN2 | THY1 | | | GRM1 | | | | CDKN1A | JUN | | | | GRIK2 | | PTGS2 | | | GRIA4 | GRM8 | | | | GRM2 |
| PSMB8 | TIE1 | | | GRM2 | | | | CHL1 | KIF3A | | | | GRIN1 | | RELA | | | GRIK2 | GTF2IRD1 | | | | GRM5 |
| PSMB9 | TNFRSF12A | | | GSK3B | | | | CHMP2B | KRAS | | | | GRIN2A | | SIRT1 | | | GRIN1 | GUCY1B3 | | | | GRM8 |
| PTGS2 | VEGFA | | | HAP1 | | | | CLDN5 | LIF | | | | GRIN2B | | SIRT2 | | | GRIN2A | HRAS | | | | GSK3B |
| SCAMP2 | XBP1 | | | HCN1 | | | | CLN8 | MAG | | | | GRIN2C | | SLC8A1 | | | GRIN2B | HTR1A | | | | HAP1 |
| SERPINB6 | | | | HDAC6 | | | | CLU | MAP2K1 | | | | GRIN2D | | SNCA | | | GRIN2C | HTR5A | | | | HOMER1 |
| SLA | | | | HIF1A | | | | CNTF | MAP2K2 | | | | GRIN3B | | SOD1 | | | GRIN2D | IGF1 | | | | HRAS |
| SLC2A1 | | | | HOMER1 | | | | CNTN1 | MAPK1 | | | | GRM1 | | SOD2 | | | GRIN3B | IGF1R | | | | ITPR3 |
| SNCA | | | | HTR5A | | | | COL4A1 | MAPK10 | | | | GRM2 | | SRC | | | GRM1 | INS | | | | KCNA1 |
| SOD2 | | | | HTT | | | | COL4A2 | MAPK3 | | | | GRM5 | | STX2 | | | GRM2 | INSR | | | | KCNB1 |
| STAMBPL1 | | | | IL1R1 | | | | COMT | MAPK8 | | | | GRM8 | | TNF | | | GRM5 | ITPR1 | | | | KCNJ10 |
| STAT1 | | | | ITPR3 | | | | CP | MAPK9 | | | | GSK3B | | TOR1A | | | GRM8 | ITPR2 | | | | KRAS |
| TGFB1 | | | | KATNA1 | | | | CR1 | MAPKAPK2 | | | | HAP1 | | TP53 | | | GSK3B | ITPR3 | | | | LAMA2 |
| TLR2 | | | | KCNA1 | | | | CREB1 | MAPT | | | | HOMER1 | | TPM1 | | | HAP1 | JUN | | | | LRRK2 |
| TMEM119 | | | | KCNB1 | | | | CREBBP | MYC | | | | HTT | | TRPM2 | | | HCN1 | KCNJ10 | | | | MAPK1 |
| TNF | | | | L1CAM | | | | CSF1R | NF1 | | | | IL1R1 | | TXNL1 | | | HOMER1 | KRAS | | | | MBP |
| TREM1 | | | | LAMA2 | | | | CTSE | NGF | | | | INSR | | UBQLN1 | | | HRAS | LPAR1 | | | | MECP2 |
| TREM2 | | | | LAMP1 | | | | CUL2 | NGFR | | | | ITGA5 | | XBP1 | | | HTR1A | MAP2K1 | | | | MPZ |
| XBP1 | | | | LPAR1 | | | | CXCL16 | NOTCH1 | | | | ITPR1 | | | | | HTR5A | MAP2K2 | | | | NCAM1 |
| | | | | LRP1 | | | | CXCR4 | NOTCH3 | | | | KCNA1 | | | | | ITPR1 | MAPK1 | | | | NF1 |
| | | | | LRRC4 | | | | CYBB | NOTCH4 | | | | KCNB1 | | | | | ITPR2 | MAPK10 | | | | NGFR |
| | | | | LRRK2 | | | | CYCS | NRG1 | | | | KCNJ10 | | | | | ITPR3 | MAPK3 | | | | NLGN4X |
| | | | | MAG | | | | DDC | NTF3 | | | | KIF3A | | | | | KCNA1 | MAPK8 | | | | NOS1 |
| | | | | MAGEE1 | | | | DDIT3 | NTNG1 | | | | LAMA2 | | | | | KCNB1 | MAPK9 | | | | NOVA1 |
| | | | | MAP2 | | | | DLG4 | NTRK1 | | | | LAMB2 | | | | | KCNJ10 | NF1 | | | | NPAS4 |
| | | | | MAPK1 | | | | DLL4 | PAK1 | | | | LAMP1 | | | | | KRAS | NFKBIA | | | | NPTN |
| | | | | MAPT | | | | DNAH1 | PDGFRB | | | | LPAR1 | | | | | MAP2K1 | NOS1 | | | | NPY |
| | | | | MBP | | | | DNAJA2 | PIK3CA | | | | LRRC4 | | | | | MAPK1 | NOS3 | | | | NQO1 |
| | | | | MTOR | | | | DRD1 | PIK3CB | | | | LRRK2 | | | | | MAPK10 | NPTN | | | | NRXN1 |
| | | | | MYH10 | | | | DRD2 | PIK3R1 | | | | MAGEE1 | | | | | MAPK3 | NPY | | | | NSF |
| | | | | NEFH | | | | EFNA1 | PLA2G16 | | | | MAPT | | | | | MAPK8 | P2RX4 | | | | NTF3 |
| | | | | NEFL | | | | EGR1 | PLA2G2A | | | | MECP2 | | | | | MAPK9 | P2RX7 | | | | NTRK1 |
| | | | | NELL2 | | | | EP300 | PLA2G2E | | | | MYD88 | | | | | NOS1 | PAK1 | | | | P2RX7 |
| | | | | NF1 | | | | EPHA2 | PLA2G2F | | | | MYH10 | | | | | NOS2 | PDE4D | | | | PARK7 |
| | | | | NLGN4X | | | | ERG | PLA2G4A | | | | NF1 | | | | | NOS3 | PIK3CA | | | | PINK1 |
| | | | | NOS1 | | | | FAS | PLA2G4B | | | | NLGN4X | | | | | NSF | PIK3CB | | | | PLA2G6 |
| | | | | NPTN | | | | FN1 | PLA2G4C | | | | NMNAT2 | | | | | P2RX4 | PIK3R1 | | | | PLCB3 |
| | | | | NRG1 | | | | FOS | PLA2G4D | | | | NOS1 | | | | | P2RX7 | PLA2G4A | | | | PLCL2 |
| | | | | NRXN1 | | | | FUS | PLA2G4E | | | | NPAS4 | | | | | PARK7 | PLA2G4B | | | | PMP22 |
| | | | | NSF | | | | FYN | PLA2G4F | | | | NPTN | | | | | PDE1B | PLA2G4C | | | | PNKD |
| | | | | NTRK1 | | | | GBA | PLCB1 | | | | NRG1 | | | | | PDGFRB | PLA2G4D | | | | PPP3CA |
| | | | | OPA1 | | | | GDNF | PLCB2 | | | | NRXN1 | | | | | PIK3CA | PLA2G4E | | | | PPP3CB |
| | | | | P2RX4 | | | | GFAP | PLCB3 | | | | NSF | | | | | PIK3CB | PLA2G4F | | | | PPT1 |
| | | | | PAK1 | | | | GJB1 | PLCB4 | | | | NTS | | | | | PIK3R1 | PLCB1 | | | | PRKACA |
| | | | | PALM | | | | GLS | PLXNB3 | | | | OLFM3 | | | | | PINK1 | PLCB2 | | | | PRKCE |
| | | | | PARK7 | | | | GNAI1 | PLXNC1 | | | | P2RX4 | | | | | PLA2G4A | PLCB3 | | | | PRKCG |
| | | | | PCSK2 | | | | GNAI2 | PPP3CA | | | | P2RX7 | | | | | PLA2G4B | PLCB4 | | | | PSEN1 |
| | | | | PINK1 | | | | GNAI3 | PPP3CB | | | | PALM | | | | | PLA2G4C | PPP2CA | | | | PTEN |
| | | | | PLCB4 | | | | GNAQ | PPP3CC | | | | PARK7 | | | | | PLA2G4D | PPP3CA | | | | PTGS2 |
| | | | | PPARGC1A | | | | GPR37 | PRKACA | | | | PFN1 | | | | | PLA2G4E | PPP3CB | | | | PTPRN2 |
| | | | | PPT1 | | | | GPR4 | PRKACB | | | | PLCB4 | | | | | PLA2G4F | PPP3CC | | | | RAB3A |
| | | | | PRKCG | | | | GRIA1 | PRKACG | | | | PPP3CA | | | | | PLCB1 | PRKACA | | | | RAC1 |
| | | | | PSEN1 | | | | GRIA2 | PRKCA | | | | PPT1 | | | | | PLCB2 | PRKACB | | | | RAPGEF2 |
| | | | | PTEN | | | | GRIN1 | PRKCB | | | | PRKACA | | | | | PLCB3 | PRKACG | | | | RIMS1 |
| | | | | PTPRN2 | | | | GRIN2A | PRKCG | | | | PRKCG | | | | | PLCB4 | PRKCA | | | | RIT2 |
| | | | | PVALB | | | | GRIN2B | PSEN1 | | | | PRKCQ | | | | | PNKD | PRKCB | | | | S100B |
| | | | | RAB3A | | | | GRIN2C | PSEN2 | | | | PSEN1 | | | | | PPP2CA | PRKCE | | | | SHANK2 |
| | | | | RET | | | | GRIN2D | PTEN | | | | PTEN | | | | | PPP2R5C | PRKCG | | | | SLC12A5 |
| | | | | SCN1A | | | | GRM5 | PTPRR | | | | PTPRN2 | | | | | PPP2R5E | PRL | | | | SLC18A2 |
| | | | | SCN2A | | | | GRN | RAC1 | | | | PVALB | | | | | PPP3CA | PTGS2 | | | | SLC18A3 |
| | | | | SHANK2 | | | | GSK3B | RAF1 | | | | RAB3A | | | | | PPP3CB | RAC1 | | | | SLC1A1 |
| | | | | SIRT2 | | | | HAP1 | RAPGEF2 | | | | RAB3C | | | | | PPP3CC | RAF1 | | | | SLC1A2 |
| | | | | SLC12A5 | | | | HDAC1 | RASGRP1 | | | | RAPGEF2 | | | | | PRKACA | RELA | | | | SLC32A1 |
| | | | | SLC18A2 | | | | HDAC2 | RELA | | | | RIMS1 | | | | | PRKACB | RHOA | | | | SLC6A3 |
| | | | | SLC1A2 | | | | HEXB | RET | | | | SHANK2 | | | | | PRKACG | RIMS1 | | | | SLC6A4 |
| | | | | SLC32A1 | | | | HGF | RHOA | | | | SLC17A6 | | | | | PRKCA | RRAS | | | | SNAP91 |
| | | | | SLC4A10 | | | | HIF1A | RRAS | | | | SLC18A2 | | | | | PRKCB | RYR1 | | | | SNCA |
| | | | | SLC6A3 | | | | HLA-DRA | RTN4 | | | | SLC18A3 | | | | | PRKCG | RYR2 | | | | SNCAIP |
| | | | | SLC8A1 | | | | HMGB1 | SHH | | | | SLC32A1 | | | | | PTGS2 | SLC17A6 | | | | SNCB |
| | | | | SLC9A6 | | | | HMOX1 | STAT1 | | | | SLC6A3 | | | | | RAF1 | SLC32A1 | | | | SORCS3 |
| | | | | SNCA | | | | HSPA6 | STAT3 | | | | SLC6A4 | | | | | RYR1 | SLC8A1 | | | | STX1A |
| | | | | SNCB | | | | HSPB1 | SYNJ1 | | | | SLC8A1 | | | | | RYR2 | SNCA | | | | STX1B |
| | | | | SOD1 | | | | HTRA2 | TGFB1 | | | | SLC9A6 | | | | | RYR3 | SOX9 | | | | STX2 |
| | | | | SPAST | | | | HTT | TGFBR2 | | | | SNCA | | | | | SCN1A | TRPV1 | | | | SYNJ1 |
| | | | | SRI | | | | ICAM2 | TNF | | | | SNCAIP | | | | | SCN2A | TSPO | | | | SYT1 |
| | | | | STAT1 | | | | IDE | TNFRSF1A | | | | SNCB | | | | | SHANK2 | | | | | SYT13 |
| | | | | SYNJ1 | | | | IDH1 | TP53 | | | | SPTBN2 | | | | | SLC12A5 | | | | | SYT4 |
| | | | | SYT1 | | | | IGF1 | | | | | SRC | | | | | SLC17A6 | | | | | SYT7 |
| | | | | SYT7 | | | | IL10 | | | | | SRI | | | | | SLC18A2 | | | | | TH |
| | | | | TENM2 | | | | IL10RA | | | | | STX1A | | | | | SLC18A3 | | | | | TNR |
| | | | | TGFB1 | | | | IL13RA1 | | | | | STX1B | | | | | SLC1A1 | | | | | TOR1A |
| | | | | TH | | | | IL15RA | | | | | STX2 | | | | | SLC1A2 | | | | | TSPO |
| | | | | THY1 | | | | IL1B | | | | | SYNJ1 | | | | | SLC32A1 | | | | | UNC13A |
| | | | | TNFRSF1B | | | | IL1R1 | | | | | SYT1 | | | | | SLC6A3 | | | | | |
| | | | | UCHL1 | | | | IL4R | | | | | SYT13 | | | | | SLC6A4 | | | | | |
| | | | | UNC13A | | | | IL6 | | | | | SYT4 | | | | | SLC8A1 | | | | | |
| | | | | WFS1 | | | | INA | | | | | SYT7 | | | | | SNCA | | | | | |
| | | | | | | | | IRF8 | | | | | TENM2 | | | | | SRC | | | | | |
| | | | | | | | | ITGA5 | | | | | TH | | | | | TH | | | | | |
| | | | | | | | | ITGAM | | | | | TOR1A | | | | | TOR1A | | | | | |
| | | | | | | | | ITPR1 | | | | | TRPV1 | | | | | TRPM2 | | | | | |
| | | | | | | | | ITPR2 | | | | | TSPO | | | | | TRPV1 | | | | | |
| | | | | | | | | ITPR3 | | | | | UNC13A | | | | | | | | | | |
| | | | | | | | | JUN | | | | | | | | | | | | | | | |
| | | | | | | | | KEAP1 | | | | | | | | | | | | | | | |

| | | | | | | | | |
|---|---|---|---|---|---|---|---|---|
| | | | | | | | | LAMB2 |
| | | | | | | | | LIF |
| | | | | | | | | LMNA |
| | | | | | | | | LRP1 |
| | | | | | | | | LRRK2 |
| | | | | | | | | LTBR |
| | | | | | | | | MAP2 |
| | | | | | | | | MAP2K1 |
| | | | | | | | | MAP2K2 |
| | | | | | | | | MAPK1 |
| | | | | | | | | MAPK10 |
| | | | | | | | | MAPK3 |
| | | | | | | | | MAPK8 |
| | | | | | | | | MAPT |
| | | | | | | | | MMP14 |
| | | | | | | | | MMP2 |
| | | | | | | | | MMP9 |
| | | | | | | | | MSN |
| | | | | | | | | MTOR |
| | | | | | | | | MYC |
| | | | | | | | | MYD88 |
| | | | | | | | | NCAM1 |
| | | | | | | | | NEFH |
| | | | | | | | | NEFL |
| | | | | | | | | NF1 |
| | | | | | | | | NFKBIA |
| | | | | | | | | NGF |
| | | | | | | | | NLRP3 |
| | | | | | | | | NMB |
| | | | | | | | | NOS1 |
| | | | | | | | | NOS2 |
| | | | | | | | | NOTCH1 |
| | | | | | | | | NPC2 |
| | | | | | | | | NPTN |
| | | | | | | | | NR4A2 |
| | | | | | | | | NTNG1 |
| | | | | | | | | OPTN |
| | | | | | | | | OSMR |
| | | | | | | | | P2RX7 |
| | | | | | | | | PALM |
| | | | | | | | | PARK7 |
| | | | | | | | | PARP1 |
| | | | | | | | | PCNA |
| | | | | | | | | PDE1B |
| | | | | | | | | PECAM1 |
| | | | | | | | | PIK3R1 |
| | | | | | | | | PINK1 |
| | | | | | | | | PLA2G4A |
| | | | | | | | | PLCB1 |
| | | | | | | | | PLCB2 |
| | | | | | | | | PLCB3 |
| | | | | | | | | PLCB4 |
| | | | | | | | | POLR2B |
| | | | | | | | | POLR2H |
| | | | | | | | | POLR2J |
| | | | | | | | | POLR2K |
| | | | | | | | | POLR2L |
| | | | | | | | | PPARGC1A |
| | | | | | | | | PPP2CA |
| | | | | | | | | PPP3CA |
| | | | | | | | | PPP3CB |
| | | | | | | | | PPP3CC |
| | | | | | | | | PRKACA |
| | | | | | | | | PRKACB |
| | | | | | | | | PRKACG |
| | | | | | | | | PRKCG |
| | | | | | | | | PRKN |
| | | | | | | | | PRNP |
| | | | | | | | | PSEN1 |
| | | | | | | | | PSEN2 |
| | | | | | | | | PSMB8 |
| | | | | | | | | PSMB9 |
| | | | | | | | | PTEN |
| | | | | | | | | PTGS2 |
| | | | | | | | | PTPRR |
| | | | | | | | | RAC1 |
| | | | | | | | | RAN |
| | | | | | | | | RELA |
| | | | | | | | | RIMS1 |
| | | | | | | | | RTN4 |
| | | | | | | | | RYR3 |
| | | | | | | | | SEC23A |
| | | | | | | | | SIRT1 |
| | | | | | | | | SIRT2 |
| | | | | | | | | SLA |
| | | | | | | | | SLC11A1 |
| | | | | | | | | SLC18A2 |
| | | | | | | | | SLC1A2 |
| | | | | | | | | SLC2A1 |
| | | | | | | | | SLC6A3 |
| | | | | | | | | SMN1 |
| | | | | | | | | SNCA |
| | | | | | | | | SNCAIP |
| | | | | | | | | SNCB |
| | | | | | | | | SOD1 |
| | | | | | | | | SOD2 |

| |
|---|
| SORL1 |
| SOX10 |
| SP1 |
| SPTBN2 |
| SQSTM1 |
| STAB1 |
| STAT3 |
| SYNJ1 |
| SYT1 |
| SYT7 |
| TAF4 |
| TAF4B |
| TARDBP |
| TBP |

| Themes | | | | | |
|---|---|---|---|---|---|
| **Compartmentalization and Structural Integrity (275)** | **Metabolism (277)** | **Neuroinflammation (134)** | **Neuron-Glia Interaction (91)** | **Neuroplasticity, Development and Aging (270)** | **Neurotransmission (278)** |
| ACHE | ABL1 | ADCY8 | ABL1 | ACVRL1 | ABAT |
| ACTN1 | ACAA1 | ADRA2A | AKT1 | ADRB2 | ACHE |
| ACVRL1 | ACHE | AMIGO1 | AKT2 | AKT1 | ADCY5 |
| ADAM10 | ACIN1 | AP3M2 | AKT3 | AKT2 | ADCY8 |
| ADCY9 | AGER | APOE | AMIGO1 | AKT3 | ADCY9 |
| ADCYAP1 | AIF1 | ARRB2 | ARHGEF10 | ANG | ADCYAP1 |
| ADORA1 | AKT1 | ATP6V0E1 | ATF4 | ANGPT2 | ADORA1 |
| ADRA2A | AKT1S1 | ATP6V1A | ATRN | APAF1 | ADORA2A |
| ADRB2 | AKT2 | ATP8A2 | BAD | APC | ADRA2A |
| AGER | AKT3 | BCL2 | BAX | ARRB2 | ADRB2 |
| AKT1 | AP1S1 | C3 | BCL2 | ATF4 | AKT1 |
| AMIGO1 | AP3M2 | CASP8 | BDNF | ATG5 | AKT2 |
| AMPH | AP3S1 | CCL2 | CALM1 | ATM | AKT3 |
| AP1S1 | AP4S1 | CCL5 | CALML5 | ATXN3 | AMPH |
| AP3M2 | APOE | CCR2 | CAMK2B | ATXN7 | AP2A2 |
| AP3S1 | APP | CCR5 | CAMK2D | BAD | AP2B1 |
| APC | ARSA | CD33 | CAMK2G | BAX | APOE |
| APOE | ATF4 | CD40 | CAMK4 | BCL2 | APP |
| APP | ATF6 | CD44 | CD9 | BCL2L1 | ARC |
| AQP4 | ATP13A2 | CD68 | CLDN5 | BDNF | ARRB2 |
| ARC | ATP6V0C | CD9 | CLU | BID | ATF4 |
| ARHGAP44 | ATP6V0D1 | CDS1 | CNTNAP1 | BRMS1L | ATP2B3 |
| ARRB2 | ATP6V1H | CLU | CTNNB1 | C3 | ATP6V0C |
| ATCAY | ATP7A | CNTF | CXCR4 | C5 | ATP6V0D1 |
| ATP6V0D1 | ATRN | CR1 | DLX1 | C6 | ATP6V0E1 |
| AVP | ATXN2 | CSF1 | DLX2 | CACNA1A | ATP6V0E2 |
| BACE1 | ATXN3 | CSF1R | EGR2 | CACNA1B | ATP6V1A |
| BCAS1 | B4GALT6 | CSF2RB | FA2H | CACNA1C | ATP6V1D |
| BCL2L1 | BAD | CSPG4 | FAM126A | CACNA1D | ATP6V1E1 |
| BECN1 | BAX | CTSE | FASLG | CACNA1F | ATP6V1G2 |
| BNIP3 | BCAS2 | CUL1 | GAL3ST1 | CACNA1S | ATP6V1H |
| CA2 | BCHE | CX3CL1 | GJB1 | CACNB2 | ATP7A |
| CACNA1A | BCL2 | CX3CR1 | GSK3B | CACNB4 | ATXN3 |
| CACNA1B | BNIP3 | CXCL10 | HEXB | CALM1 | BAD |
| CACNA1C | C9orf72 | CXCL11 | HGF | CAMK2B | BCHE |
| CACNB4 | CAB39 | CXCL12 | HRAS | CAMK2D | BCL2 |
| CADM3 | CASP3 | CXCL16 | IKBKB | CAMK2G | BDNF |
| CADPS | CAST | CXCR4 | JAM3 | CASP1 | CA2 |
| CALB1 | CCL2 | CYBB | JUN | CASP3 | CACNA1A |
| CALB2 | CCL5 | DNM1L | KCNJ10 | CASP6 | CACNA1B |
| CAMK2B | CCND1 | EGF | KEL | CASP7 | CACNA1C |
| CAMK2D | CCNH | EGFR | KLK6 | CASP8 | CACNA1D |
| CCK | CCS | EGR2 | KRAS | CASP9 | CACNA1F |
| CCL2 | CD68 | ENTPD4 | LAMA2 | CCL2 | CACNA1S |
| CCR2 | CDC27 | EPO | LPAR1 | CCL5 | CACNB2 |
| CD34 | CDC40 | FAS | MAG | CCND1 | CACNB4 |
| CD4 | CDK2 | FASLG | MAL | CCR2 | CADPS |
| CD40 | CDK5RAP3 | FLT1 | MAP2K1 | CD14 | CALB1 |
| CD8A | CDK7 | FLT4 | MAP2K2 | CD34 | CALM1 |
| CDK5 | CDS1 | FN1 | MAPK1 | CDK2 | CALML5 |
| CDK5R1 | CERS1 | GAA | MAPK10 | CDKN1A | CAMK2B |
| CHAT | CERS2 | GALC | MAPK3 | CDS1 | CAMK2D |
| CHL1 | CERS4 | GBA | MAPK8 | CHD4 | CAMK2G |
| CHRM5 | CERS6 | GNG2 | MAPK9 | CHRM5 | CAMK4 |
| CHRNA7 | CHAT | GNPTAB | MAPKAPK2 | CHRNA7 | CASP3 |
| CHRNB2 | CHMP2B | GNPTG | MBP | CNTF | CCL2 |
| CLDN15 | CLN3 | GPR84 | MOG | COL4A1 | CDK5 |
| CLDN5 | CLN8 | GPRASP1 | MTOR | COL4A2 | CHAT |
| CLN3 | CPT1B | GRN | MYRF | CREB1 | CHRM5 |
| CLN8 | CREB1 | GUSB | NF1 | CREBBP | CHRNA7 |
| CLU | CRTC2 | HGF | NFKBIA | CRTC2 | CHRNB2 |
| CNKSR2 | CTNS | HIF1A | NFKBIB | CSF1R | CNR1 |
| CNR1 | CTSE | IGF1 | NGF | CSF2RB | CNTN4 |
| CNTF | CUL1 | IL10 | NGFR | CSPG4 | COMT |
| CNTN1 | CUL2 | IL10RA | NKX6-2 | CTNNB1 | CPLX1 |
| CNTN4 | CUL3 | IL13RA1 | NRG1 | CUL1 | CREB1 |
| CNTNAP1 | CXXC1 | IL15RA | NTF3 | CUL3 | CREBBP |
| CNTNAP2 | CYBB | IL1B | NTRK1 | CX3CL1 | CRH |
| COMT | CYCS | IL1R1 | OLIG2 | CX3CR1 | CTNNB1 |
| CPLX1 | DDIT3 | IL4R | PDPK1 | CXCL10 | CX3CL1 |
| CREB1 | DDX23 | IL6 | PIK3CA | CXXC1 | CX3CR1 |
| CRH | DGKB | IL6R | PIK3CB | CYCS | CYP4X1 |
| CTNNB1 | DGKE | INHBB | PIK3R1 | DDIT3 | DAGLA |
| CX3CL1 | DLAT | IRF8 | PLLP | DLL4 | DBH |
| DAGLA | DLD | ITGA5 | PMP22 | DOT1L | DDC |
| DCX | DNAJA2 | ITGAL | PSEN1 | DRD2 | DLG3 |

| | | | | | |
|---|---|---|---|---|---|
| DDC | DNM2 | ITGAX | PSEN2 | EFNA1 | DLG4 |
| DES | EIF2S1 | JAM3 | PTEN | EFNA5 | DLGAP1 |
| DLG3 | ENTPD4 | LIF | RAC1 | EFNB3 | DNM2 |
| DLG4 | EP300 | LMNA | RAF1 | EFR3A | DRD1 |
| DLGAP1 | ERLEC1 | LOX | RELA | EGF | DRD2 |
| DNAH1 | FAS | LTBR | RHOA | EGFL7 | DRD4 |
| DNM1L | FN1 | MARCO | SCN2A | EGFR | EGFR |
| DNM2 | FOS | MMP12 | SH3TC2 | EGR1 | EGR2 |
| DRD1 | FXN | MMP14 | SIRT2 | EHMT1 | ENTPD2 |
| DRD2 | GAA | MMP16 | SOD1 | EIF2S1 | EP300 |
| DRD4 | GAL3ST1 | MMP19 | TGFB1 | EMCN | ERBB3 |
| EFNA1 | GALC | MMP2 | TP53 | EMP2 | F2 |
| EFNA5 | GBA | MMP24 | UGT8 | ENG | FGF12 |
| EFNB3 | GDPD2 | MMP9 | XK | EP300 | FGF14 |
| EPHA2 | GFPT1 | NAGLU | ZNF24 | EPHA2 | FMR1 |
| EPHA3 | GGA1 | NGFR | | EPHA3 | FOS |
| EPHA4 | GGT1 | NLRP3 | | EPHA4 | FYN |
| EPHA5 | GLS | NPAS4 | | EPHA5 | GABRA1 |
| EPHA6 | GNAO1 | NPC2 | | EPHA6 | GABRA4 |
| EPHA7 | GNPTAB | OPTN | | ERBB3 | GABRB2 |
| ESAM | GNPTG | OSMR | | ERG | GABRB3 |
| FAS | GPD1L | P2RX4 | | FAM126A | GABRG2 |
| FMR1 | GPR37 | P2RY12 | | FAS | GABRP |
| FRMPD4 | GSK3B | PDGFRB | | FASLG | GABRR1 |
| FUS | GSR | PGK1 | | FGF12 | GABRR3 |
| FYN | GSS | PLCL2 | | FGF14 | GAD1 |
| GABRA1 | GSTP1 | PLEKHO2 | | FGF2 | GAD2 |
| GABRA4 | GTF2A1 | PLXNC1 | | FLT1 | GDNF |
| GABRB2 | GTF2B | PPM1L | | FLT4 | GFAP |
| GABRB3 | GTF2H1 | PRL | | FMR1 | GLRB |
| GABRG2 | GTF2H3 | PSEN2 | | FN1 | GLS |
| GABRP | GTF2IRD1 | PSMB8 | | FOS | GNAI1 |
| GABRR1 | GUCY1B3 | PSMB9 | | FYN | GNAI2 |
| GABRR3 | GUSB | PTGS2 | | GATA2 | GNAI3 |
| GAD1 | HDAC2 | SCAMP2 | | GDNF | GNAO1 |
| GAD2 | HDAC6 | SERPINB6 | | GPR4 | GNAQ |
| GFAP | HEXB | SLA | | GSK3B | GNB5 |
| GLRB | HGF | SLC2A1 | | HDAC1 | GNG2 |
| GNAI2 | HIF1A | SNCA | | HDAC2 | GNGT1 |
| GNAO1 | HMOX1 | SOD2 | | HDAC6 | GPR37 |
| GNAQ | HNRNPM | STAMBPL1 | | HDAC7 | GRIA1 |
| GRIA1 | HPGDS | STAT1 | | HGF | GRIA2 |
| GRIA2 | HSPA6 | TGFB1 | | HIF1A | GRIA3 |
| GRIA3 | HSPB1 | TGFBR2 | | HMGB1 | GRIA4 |
| GRIA4 | HTRA2 | TLR2 | | HMOX1 | GRIK2 |
| GRIK2 | IDE | TMEM119 | | HRAS | GRIN1 |
| GRIN1 | IDH1 | TNF | | HSPA6 | GRIN2A |
| GRIN2A | IGF1 | TNFRSF10A | | HSPB1 | GRIN2B |
| GRIN2B | IGF1R | TNFRSF10B | | HTRA2 | GRIN2C |
| GRIN2C | IL1B | TNFRSF10D | | IGF1 | GRIN2D |
| GRIN2D | IL1R1 | TNFRSF11B | | IKBKB | GRIN3B |
| GRIN3B | IL6 | TNFRSF12A | | IL1B | GRM1 |
| GRM1 | INS | TNFRSF1A | | IL1R1 | GRM2 |
| GRM2 | INSR | TNFRSF1B | | IL6 | GRM5 |
| GRM5 | IPCEF1 | TREM1 | | INPP4A | GRM8 |
| GRM8 | JUN | TREM2 | | INPP5F | GSK3B |
| GSK3B | KEAP1 | VEGFA | | ITGA5 | GTF2IRD1 |
| GSN | LAMP1 | XBP1 | | ITPR1 | GUCY1B3 |
| HAP1 | LCLAT1 | | | ITPR2 | HAP1 |
| HCN1 | LDHC | | | ITPR3 | HCN1 |
| HDAC6 | LMNA | | | JAM3 | HOMER1 |
| HIF1A | LPO | | | JUN | HRAS |
| HLA-DRA | LRRK2 | | | KIF3A | HTR1A |
| HOMER1 | LSM2 | | | KRAS | HTR5A |
| HTR5A | LSM7 | | | LIF | IGF1 |
| HTT | LSR | | | LMNA | IGF1R |
| ICAM1 | LYPLA1 | | | MAG | INS |
| ICAM2 | MAN2B1 | | | MAP2K1 | INSR |
| IL1R1 | MAPK10 | | | MAP2K2 | ITPR1 |
| INA | MAPK8 | | | MAPK1 | ITPR2 |
| INSR | MAPK9 | | | MAPK10 | ITPR3 |
| ISLR2 | MAPT | | | MAPK3 | JUN |
| ITGA5 | MFN2 | | | MAPK8 | KCNA1 |
| ITGA7 | MGMT | | | MAPK9 | KCNB1 |
| ITGAL | MMP14 | | | MAPKAPK2 | KCNJ10 |
| ITGAM | MNAT1 | | | MAPT | KRAS |
| ITPR1 | MTOR | | | MEAF6 | LAMA2 |
| ITPR3 | MUTYH | | | MECP2 | LPAR1 |
| JAM3 | NAGLU | | | MMP14 | LRRK2 |
| KATNA1 | NAPSA | | | MMP19 | MAP2K1 |
| KCNA1 | NCF1 | | | MMP2 | MAP2K2 |
| KCNB1 | NEFH | | | MMRN2 | MAPK1 |
| KCNJ10 | NFE2L2 | | | MTA1 | MAPK10 |
| KIF3A | NGFR | | | MTA2 | MAPK3 |
| L1CAM | NME5 | | | MTHFR | MAPK8 |
| LAMA2 | NOL3 | | | MYC | MAPK9 |
| LAMB2 | NOS1 | | | NCL | MBP |
| LAMP1 | NOS3 | | | NELFA | MECP2 |
| LPAR1 | NPC1 | | | NF1 | MPZ |
| LRP1 | NPC2 | | | NFKBIA | NCAM1 |
| LRRC4 | NPY | | | NGF | NF1 |
| LRRK2 | NQO1 | | | NGFR | NFKBIA |
| MAG | NR4A2 | | | NOS1 | NGFR |
| MAGEE1 | OXR1 | | | NOS3 | NLGN4X |
| MAP2 | PARK7 | | | NOTCH1 | NOS1 |
| MAPK1 | PARP1 | | | NOTCH3 | NOS2 |

| | | | | | |
|---|---|---|---|---|---|
| MAPT | PCNA | | | NOTCH4 | NOS3 |
| MBP | PDGFRB | | | NRG1 | NOVA1 |
| MECP2 | PDPK1 | | | NRXN1 | NPAS4 |
| MPZ | PGAM1 | | | NTF3 | NPTN |
| MSN | PGK1 | | | NTNG1 | NPY |
| MTOR | PIK3CA | | | NTRK1 | NQO1 |
| MYD88 | PIK3CB | | | PAK1 | NR4A2 |
| MYH10 | PIK3R1 | | | PARP1 | NRXN1 |
| NCAM1 | PINK1 | | | PCNA | NSF |
| NEFH | PLA2G16 | | | PDGFRB | NTF3 |
| NEFL | PLA2G2A | | | PDPK1 | NTRK1 |
| NEGR1 | PLA2G2E | | | PHF19 | P2RX4 |
| NELL2 | PLA2G2F | | | PHF2 | P2RX7 |
| NEO1 | PLA2G4A | | | PHF21A | PAH |
| NES | PLA2G4B | | | PIK3CA | PAK1 |
| NF1 | PLA2G4C | | | PIK3CB | PARK7 |
| NINJ2 | PLA2G4D | | | PIK3R1 | PDE1B |
| NLGN4X | PLA2G4E | | | PKN1 | PDE4D |
| NMNAT2 | PLA2G4F | | | PLA2G16 | PDGFRB |
| NOS1 | PLA2G6 | | | PLA2G2A | PIK3CA |
| NPAS4 | POLR2B | | | PLA2G2E | PIK3CB |
| NPTN | POLR2H | | | PLA2G2F | PIK3R1 |
| NRG1 | POLR2J | | | PLA2G4A | PINK1 |
| NRXN1 | POLR2K | | | PLA2G4B | PLA2G4A |
| NSF | POLR2L | | | PLA2G4C | PLA2G4B |
| NTNG1 | PPARGC1A | | | PLA2G4D | PLA2G4C |
| NTRK1 | PPP2CA | | | PLA2G4E | PLA2G4D |
| NTS | PPP2R5C | | | PLA2G4F | PLA2G4E |
| OLFM3 | PPP2R5E | | | PLCB1 | PLA2G4F |
| OPA1 | PPT1 | | | PLCB2 | PLA2G6 |
| P2RX4 | PQBP1 | | | PLCB3 | PLCB1 |
| P2RX7 | PRKAA2 | | | PLCB4 | PLCB2 |
| PAK1 | PRKCSH | | | PLXNB3 | PLCB3 |
| PALM | PRKN | | | PLXNC1 | PLCB4 |
| PARK7 | PRNP | | | PPARGC1A | PLCL2 |
| PCSK2 | PRPF3 | | | PPP3CA | PMP22 |
| PECAM1 | PRPF31 | | | PPP3CB | PNKD |
| PFN1 | PSEN1 | | | PPP3CC | PPP2CA |
| PINK1 | PSMB8 | | | PRF1 | PPP2R5C |
| PLCB4 | PSMB9 | | | PRKAA2 | PPP2R5E |
| PLS1 | PTDSS1 | | | PRKACA | PPP3CA |
| PPARGC1A | PTDSS2 | | | PRKACB | PPP3CB |
| PPP3CA | PTGS2 | | | PRKACG | PPP3CC |
| PPT1 | RAB2A | | | PRKCA | PPT1 |
| PRKACA | RAD23B | | | PRKCB | PRKACA |
| PRKCG | RAN | | | PRKCG | PRKACB |
| PRKCQ | RELA | | | PSEN1 | PRKACG |
| PSEN1 | SART1 | | | PSEN2 | PRKCA |
| PTEN | SEC23A | | | PTEN | PRKCB |
| PTPRN2 | SF3A2 | | | PTGS2 | PRKCE |
| PVALB | SF3B2 | | | PTPRR | PRKCG |
| RAB3A | SF3B4 | | | RAC1 | PRL |
| RAB3C | SGPL1 | | | RAF1 | PSEN1 |
| RAPGEF2 | SIRT1 | | | RAPGEF2 | PTEN |
| RDX | SIRT2 | | | RASGRP1 | PTGS2 |
| RET | SLC11A1 | | | RELA | PTPRN2 |
| RIMS1 | SLC1A1 | | | RET | RAB3A |
| SCN1A | SLC1A2 | | | RHOA | RAC1 |
| SCN2A | SLC8A1 | | | RING1 | RAF1 |
| SHANK2 | SLU7 | | | RRAS | RAPGEF2 |
| SIRT2 | SMN1 | | | RTN4 | RELA |
| SLC12A5 | SMPD4 | | | SHH | RHOA |
| SLC17A6 | SNCA | | | SIRT1 | RIMS1 |
| SLC18A2 | SNRPA | | | SIRT2 | RIT2 |
| SLC18A3 | SOD1 | | | SIRT7 | RRAS |
| SLC1A2 | SOD2 | | | SMYD1 | RYR1 |
| SLC2A1 | SQSTM1 | | | SNCA | RYR2 |
| SLC32A1 | SRC | | | SOX10 | RYR3 |
| SLC4A10 | SRSF4 | | | SP1 | S100B |
| SLC6A3 | STX2 | | | SP100 | SCN1A |
| SLC6A4 | SUCLA2 | | | SPI1 | SCN2A |
| SLC8A1 | TAF10 | | | STAB1 | SHANK2 |
| SLC9A6 | TAF4 | | | STAT1 | SLC12A5 |
| SNCA | TAF4B | | | STAT3 | SLC17A6 |
| SNCAIP | TAF6L | | | SYNJ1 | SLC18A2 |
| SNCB | TAF9 | | | TAF10 | SLC18A3 |
| SOD1 | TARDBP | | | TAF6L | SLC1A1 |
| SPAST | TAZ | | | TAF9 | SLC1A2 |
| SPTBN2 | TBP | | | TGFB1 | SLC32A1 |
| SRC | TBPL1 | | | TGFBR2 | SLC6A3 |
| SRI | TCERG1 | | | THY1 | SLC6A4 |
| STAT1 | TCIRG1 | | | TIE1 | SLC8A1 |
| STX1A | TFAM | | | TLR4 | SNAP91 |
| STX1B | TNF | | | TNF | SNCA |
| STX2 | TOR1A | | | TNFRSF10A | SNCAIP |
| SYNJ1 | TP53 | | | TNFRSF10B | SNCB |
| SYT1 | TPM1 | | | TNFRSF10D | SORCS3 |
| SYT13 | TRIM37 | | | TNFRSF12A | SOX9 |
| SYT4 | TRPM2 | | | TNFRSF1A | SRC |
| SYT7 | TWISTNB | | | TP53 | STX1A |
| TENM2 | TXNL1 | | | TRADD | STX1B |
| TGFB1 | U2AF2 | | | TRIM28 | STX2 |
| TH | UBE2K | | | TRIM37 | SYNJ1 |
| THY1 | UBE2N | | | UBE2N | SYT1 |
| TNC | UBE3A | | | USP21 | SYT13 |
| TNFRSF1B | UBQLN1 | | | VEGFA | SYT4 |

| | | | | | |
|---|---|---|---|---|---|
| TOR1A | UGCG | | | XBP1 | SYT7 |
| TPM1 | UGT8 | | | XIAP | TCIRG1 |
| TRPV1 | USP30 | | | | TH |
| TSPO | VCP | | | | TNR |
| UCHL1 | VIP | | | | TOR1A |
| UNC13A | WFS1 | | | | TRPM2 |
| WFS1 | XAB2 | | | | TRPV1 |
| | XBP1 | | | | TSPO |
| | XIAP | | | | UNC13A |
| | | | | | VIP |
| **Reference genes** | | | | | |
| **(10)** | | | | | |
| AARS | | | | | |
| ASB7 | | | | | |
| CCDC127 | | | | | |
| CNOT10 | | | | | |
| CSNK2A2 | | | | | |
| FAM104A | | | | | |
| LARS | | | | | |
| MTO1 | | | | | |
| SUPT7L | | | | | |
| TADA2B | | | | | |
